# ERBB signalling contributes to immune evasion in KRAS-driven lung adenocarcinoma

**DOI:** 10.1101/2023.07.24.550274

**Authors:** Sarah Laing, Björn Kruspig, Robin Shaw, Leah Officer-Jones, Sarah Edwards, Danielle McKinven, Ya-Ching Hsieh, Ian Powley, Nicola Brady, Rachel Pennie, Ryan Kwan, Anthony Lima, Szymon Myrta, Manikandan Periyasamy, Isabel C Dye, Colin Nixon, Graeme Clark, Melissa R. Junttila, Danilo Maddalo, Crispin Miller, Simak Ali, Matthew J. Fuchter, Dorothee Nickles, Kristina Kirschner, Robert B. Brown, John Le Quesne, Douglas Strathdee, Seth B. Coffelt, Ed Roberts, Daniel J. Murphy

**Author notes:** **Conflict of Interest Statement**. DJM has received funding from Puma Biotechnology to investigate use of Neratinib in KRAS-mutant lung cancer mouse models. DJM has additionally received funding from the Merck Group for work unrelated to the present subject. DN, DM & AL are employees of Genentech Inc. SM is employed by Roche Global IT Solutions Centre. MJ is an employee and stock-holder of ORIC Pharmaceuticals. RBB and MJF are inventors on a patent including HKMTI-1-005. All other authors declare no financial conflicts of interest.

## Abstract

Immunotherapy is increasingly viewed as treatment of choice for lung cancer, however, clinical responses to immune checkpoint blockade remain highly unpredictable and are largely transient. A deeper mechanistic understanding of the dynamics of tumour:immune interactions is needed to drive rational development of improved treatment strategies. Progress is hampered by a paucity of autochthonous model systems in which to interrogate the 2-way interactions of immune responses to evolving tumours and vice-versa. Specifically, commonly used genetically engineered mouse models typically lack the genetic diversity needed to drive an adaptive immune response. APOBEC mutagenesis signatures are prominent in lung cancer and APOBEC activity is predicted to drive immune visibility through Cytidine deaminase activity, coupled with inaccurate DNA-repair responses. We therefore generated a CRE-inducible *APOBEC3B* allele, interbred with multiple oncogenic drivers of lung adenocarcinoma, and used the resulting mice to investigate the response to PD1 blockade at single cell resolution.

SIGNIFICANCE

Using our novel immune-visible model of KRas-driven autochthonous lung adenocarcinoma, we uncovered a surprising increase in tumour-cell expression of EGFR/ERBB ligands following treatment with α-PD1 and present evidence that transient ERBB blockade can restore immune surveillance in KRas mutant LuAd and combine effectively with immune checkpoint blockade.

## INTRODUCTION

Immune checkpoint blockade (ICB) has transformed clinical oncology and is now widely approved for front-line treatment of many cancer types, including cancers of the lung and thorax (1–4). Targeted blockade of the canonical immune checkpoint pairing of CTLA-4 and PD1/PD-L1, or combining PD1/PD-L1 blockade with conventional treatment modalities, has yielded remarkable outcomes in substantial subsets of cancer patients, yet immediate responses are highly variable and durable benefit remains infrequent, particularly for lung and thoracic cancer patients (5–8). A deeper understanding of the dynamic co-evolution of tumour and anti-tumour immune responses is needed to identify new strategies to increase both frequency and duration of benefit from ICB therapy. However, there is currently a paucity of fully immunocompetent *in vivo* model systems in which to investigate these dynamics: syngeneic models rely on transplantation of large numbers of fully transformed tumour cells that deny the opportunity for studying early tumour evolution and are known to provoke inflammatory responses that can skew subsequent immune interactions (9–11); humanised models, such as those using transplantation of human hematopoietic stem cells, are limited by their short duration and complicated by their background of graft-versus-host disease (12); sporadically inducible genetically engineered mouse (GEM) models allow for autochthonous co-evolution of tumour and histocompatible immune responses, but lack the genetic complexity of human tumours and are typically immune-silent (13, 14). Recent efforts to generate immune-visible GEM models have partially addressed this latter point (15–17), however their reliance on individual strongly antigenic epitopes may not fully reflect the polyclonal, multi immune effector lineage responses to a spectrum of neoantigens.

APOBEC Cytidine deaminases are endogenous single-stranded nucleic acid-modifying enzymes that play major roles in cell-intrinsic anti-viral responses and suppress expression of endogenous retroviruses and transposons (18, 19). APOBEC mutagenesis signatures are prominent in many cancer types and APOBEC3 enzymes are thought to contribute substantially to genetic heterogeneity and tumour evolution in lung cancer (20–23). The alteration of coding sequences or aberrant expression of occult transcripts arising from APOBEC mutagenesis in cancer might therefore be expected to generate neoantigens that provoke adaptive immune responses while simultaneously driving tumour evolution (24–26). With this in mind, we generated a novel murine allele for conditional expression of APOBEC3B from the *Rosa26* locus and interbred the resulting mice with well-established allelic combinations known to yield progressive lung adenocarcinoma (LuAd) (27–30). Here we report the generation of a novel immune-visible GEMM of KRAS-driven autochthonous LuAd and explore the dynamics of the tumour response to PD1 blockade, alone and in combination with targeted therapeutics. Our data suggest that KRAS mutant lung tumours progress to more aggressive phenotypes under the selective pressure of an effective immune response, at least in part via upregulation of ERBB signalling. Targeted suppression of ERBB may thus enhance KRAS-mutant LuAd responses to ICB therapy.

## RESULTS

### Co-expression of APOBEC3B extends survival of a subset of LuAd mouse models

To model the impact of APOBEC3B (henceforth A3B) on lung cancer development we generated a conditional allele enabling CRE-dependent expression of A3B from the endogenous *Rosa26* locus, *Rosa26^DS-lsl-APOBEC3B^*, and interbred the resulting mice with various oncogenic allelic combinations known to drive progressive lung adenocarcinoma. Combinations of conditional alleles were activated in adult mice by intranasal installation of replication defective, lineage-restricted, Adeno-SPC-CRE virus (AS-CRE) and mice were maintained until clinical signs required humane killing. Inclusion of the A3B allele (KMA) clearly extended survival of mice bearing lung tumours driven by endogenously expressed KRAS^G12D^ and modestly overexpressed c-MYC (KM), the latter also expressed conditionally from the *Rosa26* locus (Fig. 1A, B). Comparison of KM with KMA lungs at clinical endpoint revealed no difference in overall tumour burden or histological appearance, however, metastases to the liver, identified by CRE-dependent expression of an IRFP reporter allele, were found in a subset of KMA but no KM mice (Fig. 1 C, D & S1 A). Analysis of the lungs of pre-symptomatic mice, harvested at 8 weeks post induction (p.i.), showed significantly lower tumour burden in KMA compared with KM mice. BrdU staining revealed no difference in tumour proliferation at this time while cleaved Caspase 3 tended to be higher in KMA tumours (Fig. 1E, F). A3B was strongly expressed in tumours of all KMA mice analysed at either 8 weeks p.i. or at endpoint and was entirely absent from both KM tumours and non-transformed KMA lung tissue (Fig. S1B). Immunoblotting of lysates from lungs of tumour bearing end-stage mice revealed no consistent differences in expression of KRAS^G12D^ or MYC (Fig. S1C).

**Figure 1:**
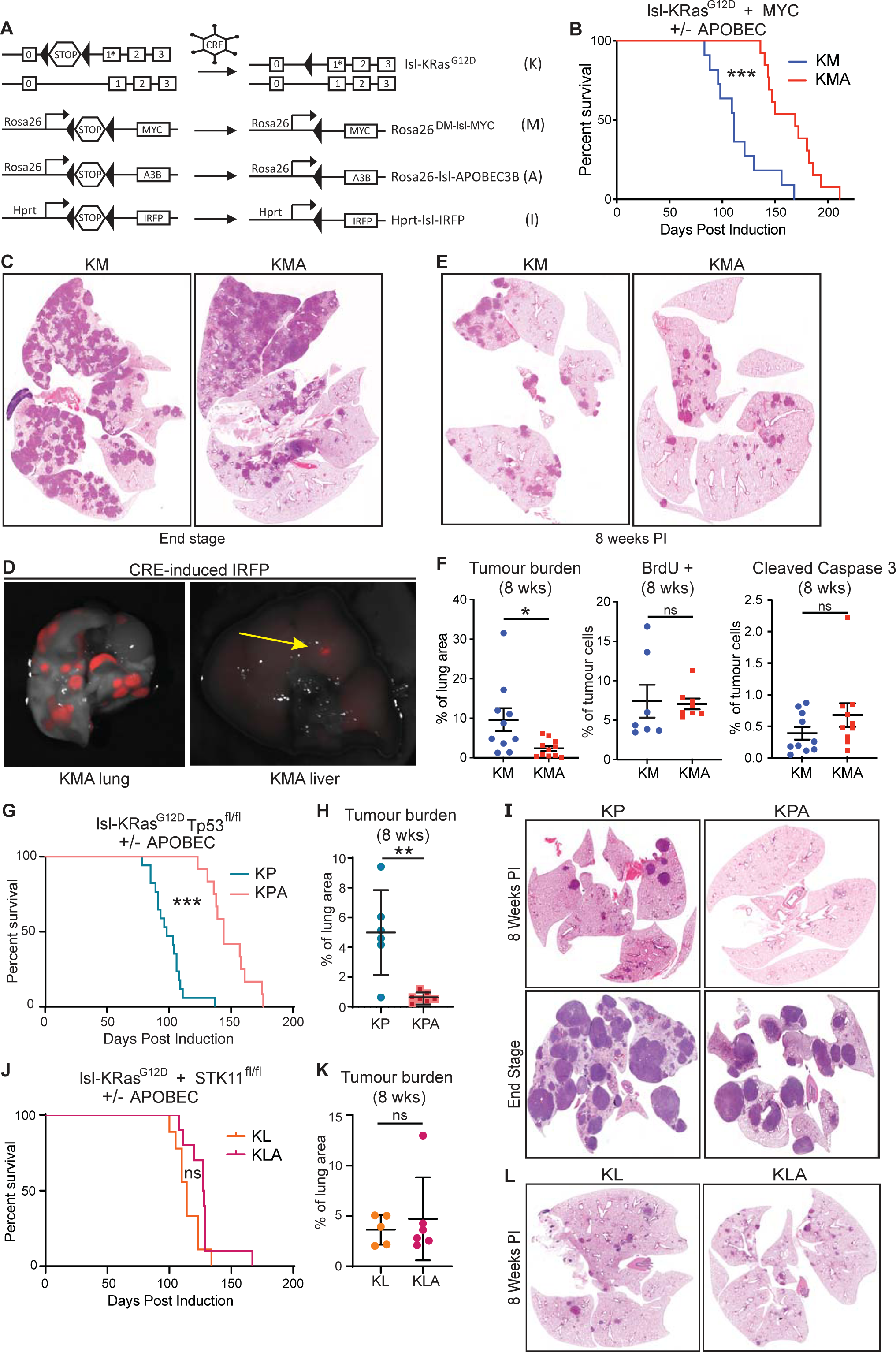
Context dependent survival extension in model of LuAd upon co-expression of APOBEC3B. **A)** Schematic of alleles used in A-F and Figures 2-7. **B)** Overall survival of KM (N = 11) versus KMA (N = 13) mice, measured in days after allele induction with 1x10^8 pfu AS-CRE. Mantel-Cox log rank test. **C)** Representative images show histological tumour burden [haematoxylin and eosin (H&E)] of lungs from KM and KMA mice at end-stage. **D)** Detection of IRFP expression (from Hprt-lsl-IRFP) in primary lung tumours (left) and a liver metastasis (right) in a KMA mouse induced at a viral titre of 1X10^7 pfu SPC-Cre, harvested at end-stage. **E)** Representative images show H&E stained lung tumour burden of KM and KMA mice at 8-weeks post allele induction. **F)** Left panel: overall lung tumour burden calculated as a percentage of total lung area at 8-weeks post allele induction in KM (N = 10) and KMA (N = 11) mice. Centre panel: quantification of percentage of BrdU IHC cells in lung tumours in KM (N = 7) and KMA (N = 8) mice at 8-weeks post allele induction. Right panel: quantification of cleaved Caspase3 IHC in KM (N=10) and KMA (N=9) at 8 weeks post induction. Each data point represents the average score of all tumours from a section in an individual mouse. Error bars represent Mean ± SEM (unpaired t-test). **G)** Overall survival of KP (N = 17) versus KPA (N = 13) mice measured in days post allele induction with 1x10^8 AS-CRE. Mantel-Cox log rank test. **H)** Overall lung tumour burden calculated as a percentage of total lung area at 8-weeks post allele induction in KP (N = 6) and KPA (N = 7) mice. Error bars represent Mean ± SD (unpaired t-test). **I)** Representative images show H&E stained tumour burden of KP and KPA mice at 8-weeks post allele induction and at end-stage. **J)** Overall survival of KL (N = 9) and KLA (N = 10) mice measured in days post allele induction with 1x10^8 AS-CRE. Mantel-Cox log rank test. **K)** Overall lung tumour burden calculated as a percentage of total lung area at 8-weeks post allele induction in KL (N = 5) and KLA (N = 6) mice. Error bars represent Mean ± SD. **L)** Representative images show tumour burden histology, H&E, of KL and KLA mice at 8-weeks post allele induction. For all panels, ns denotes not significant; * denotes P <0.05; ** denotes P <0.01; *** denotes P < 0.001.

*Rosa26^DS-lsl-APOBEC3B^* mice were additionally interbred with mice carrying *LSL-KRas^G12D^*combined with homozygous floxed alleles of either *Trp53* (KP; KPA)(31) or *Stk11,* encoding LKB1 (KL; KLA) (32). As found for KMA mice, KPA mice showed markedly increased survival over KP comparators (Fig. 1G). Analysis of lungs harvested at 8 weeks p.i. revealed a profound suppression of tumour development in KPA lungs, however, endpoint tumour burden was equivalent in KPA and KP mice (Fig. 1H, I). In sharp contrast, we found no difference in survival between KLA and KL mice, and tumour burden at 8 weeks p.i. was indistinguishable (Fig. 1J-L).

### Weak activity of APOBEC3B Cytidine deaminase expressed from *Rosa26*

Mutation signatures linked to inferred APOBEC activity are prominent in human LuAd, although the timing, mechanism and extent of APOBEC-driven mutagenesis remain active areas of investigation (22, 33–35). In the context of DNA, the Cytidine deaminase activity of A3B converts Cytidine to Uridine, prompting activation of DNA-damage response pathways to replace the disallowed nucleotide, typically with Thymidine or Guanosine (24). We therefore examined our murine A3B-expressing tumours for evidence of elevated mutagenesis. Whole exome sequencing of end-stage tumours failed to reveal consistent elevation of mutation burden in KMA versus KM tumours. COSMIC signature analysis (36) moreover did not identify canonical APOBEC mutagenesis signatures (Figure S2A, B). To directly measure A3B activity, we generated cell lines from end-stage KPA tumours (repeated attempts to do so from KM/KMA tumours were unsuccessful) and measured *in vitro* Cytidine deaminase activity in cell lysates. Consistent with strong expression of A3B protein in KPA-derived cell lines, KPA lysates showed significantly elevated Cytidine deaminase activity compared with KP lysates (Fig S2C, D). An independent assay to quantify abasic DNA sites similarly showed increased DNA damage in KPA cells compared with KP cells (Fig S2E).

### APOBEC3B drives a transient immune response in KM lung tumours

The increased survival of KMA over KM mice, and indeed the difference in tumour burden at 8 weeks p.i., was not explained by differential rates of proliferation or by differential expression of driver oncogenes. We therefore harvested tumours at 8 weeks p.i. and performed bulk tumour RNA-SEQ analysis. Surprisingly, the vast majority of transcripts showed no significant difference between KMA and KM tumours (Fig. 2A) indicating that A3B expression does not strongly alter the early tumour phenotype. Metacore GeneGO pathway analysis however revealed enrichment of immune-related gene expression, particularly type I and type II Interferon pathway activity, in KMA tumours (Fig. 2A). Analysis of individual genes showed significantly increased expression of Interferon inducible genes *Ifitm1, Ifitm3*, and multiple oligoadenylate synthase (*Oas*) genes, which all contribute to Interferon-induced antiviral activity; high expression of *IghM*, suggesting infiltration of naïve B cells; and high expression of the dendritic and effector immune cell chemokine *Ccl20* (Fig. S3 A-C). Notably, the tumour sample showing highest expression of *Ccl20* also showed highest expression of T cell-related genes, along with the T cell chemoattractant pairing of *Cxcl10* and *Cxcr3* (Fig. S3D). We therefore characterised the tumour immune microenvironment by IHC. Staining revealed strikingly higher infiltration of KMA tumours by CD8 and CD4 T cells, as compared with KM tumours, at the 8-week timepoint (Fig. 2B). 8-week p.i. KMA tumours additionally showed increased NK (NKp46) and B cell (B220/CD45R) infiltration, while tumour infiltration by Macrophages (F4/80) and Neutrophils (Ly6G) was equivalent between the 2 genotypes (Fig. S4A, B). CD45R staining revealed increased frequency of tumour-proximal tertiary lymphoid structures in KMA lungs, and TLS number correlated with T cell infiltration (Fig. S4C, D). Analysis of lungs from end-stage mice showed near complete exclusion of T cells from both KM and KMA tumours along with no obvious difference in other immune populations (Fig. 2B-D, S4E). We therefore examined a cohort of mice at 12 weeks p.i.. Remarkably, this intermediate time point also showed near complete exclusion of T cells from tumours of both genotypes, revealing the sharply transient nature of T cell tumour infiltration during early tumour development in KMA mice (Fig. 2B-D).

**Figure 2:**
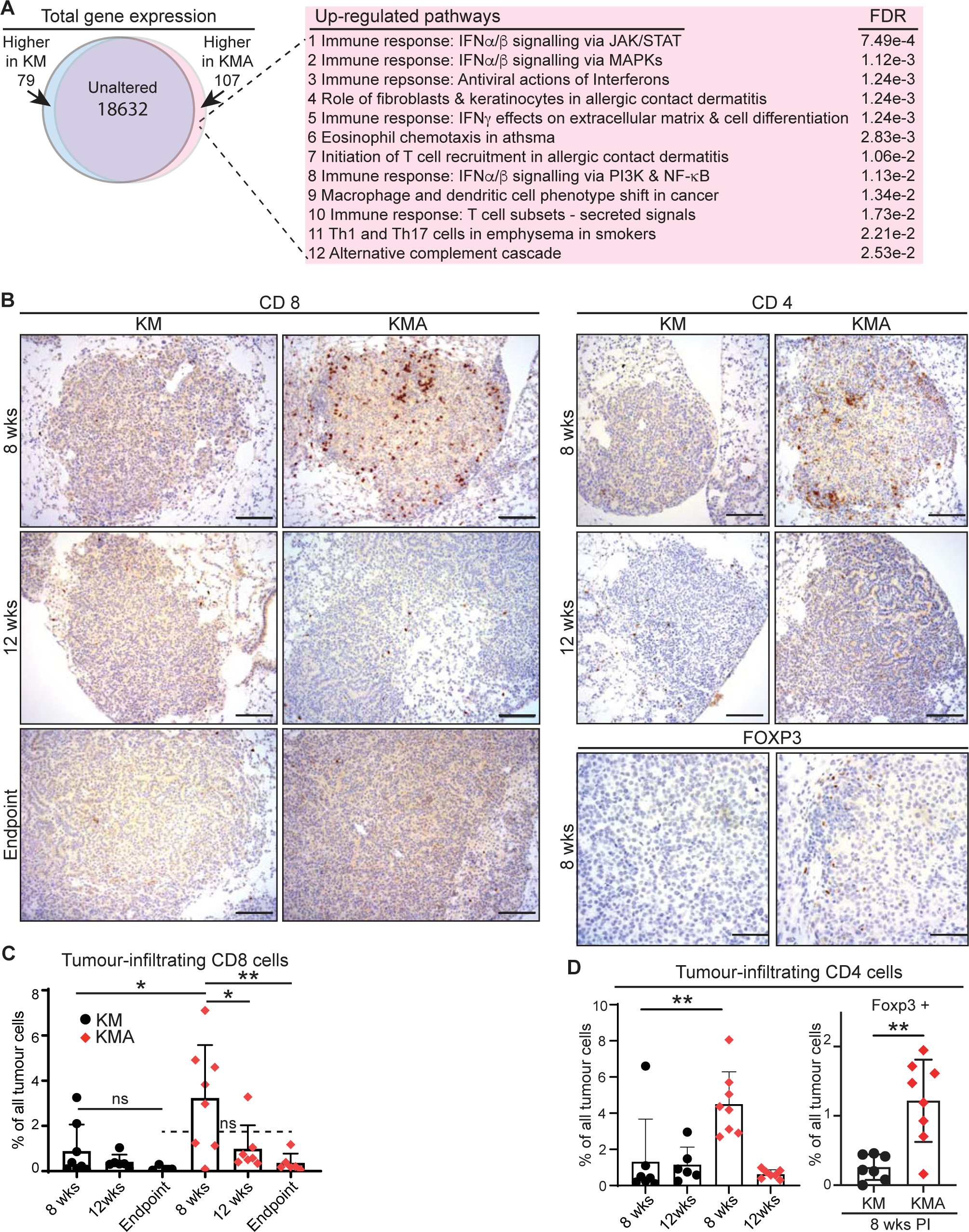
Expression of A3B results in a transient T cell response to LuAd. **A)** Schematic of gene expression changes from tumour bearing lungs of KM and KMA (N = 4 biological replicates) mice taken at 8 weeks post allele induction and analysed by RNA-Seq. The top 12 significantly modulated pathways in KMA mice were identified using Metacore GeneGO analysis. **B)** Representative images of CD8α stained KM and KMA tissue sections harvested at 8- and 12-weeks post allele induction and from end-stage animals, CD4 stained tissue sections at 8- and 12-weeks post allele induction, and FoxP3+ stained tissue sections at 8-weeks post allele induction, illustrating T cell infiltration into KM and KMA tumours throughout tumour progression. Scale bars, 100µm. **C)** QuPath quantification of the percentage of cells in tumours that are CD8α+ T cells in KM and KMA mice at 8- (N = 8 of each genotype) and 12-weeks (N = 6 KM & 7 KMA) post allele induction and from end-stage mice (N = 4 KM & 6 KMA). Each data point represents the average score of all tumours from a section in an individual mouse. Error bars represent Mean ± SD (unpaired t-test). **D)** QuPath quantification of the percentage of cells in tumours that are CD4+ T cells in KM and KMA mice at 8- (N = 7 KM and 8 KMA) and 12-weeks (N = 6 KM and 7 KMA) post allele induction. QuPath quantification of tumour-infiltrating FoxP3+ cells in KM and KMA mice at 8-weeks post allele induction (N = 7 KM and 8 KMA). Each data point represents the average score of all tumours in a section from an individual mouse. Error bars represent Mean ± SD (unpaired t-test). For all panels, ns denotes not significant; * denotes P <0.05; ** denotes P <0.01.

### Depletion of CD8 T cells negates the survival advantage of KMA mice

We asked if the tumour-infiltrating effector T cells might account for the survival advantage seen in KMA mice over KM counterparts. CD8 T cells were transiently depleted from KM and KMA mice by 3 weeks treatment with CD8-depleting antibody (clone 2.43), commencing from day 50 p.i., ie. during the active phase of KMA tumour infiltration (Fig. 3A). Flow cytometry from blood samples taken after 2 weeks of treatment confirmed systemic depletion of CD8 T cells (Fig. S5A, B). Depletion of CD8 T cells significantly reduced survival of KMA mice but had no such effect on KM counterparts (Fig. 3B, C). Indeed, transient depletion of CD8 T cells completely negated the survival difference between KM and KMA mice (Fig. S5C). We also used a genetic approach to T cell suppression by interbreeding both KM and KMA mice with constitutive *Tcrb* knockout mice, deficient for both CD4 and CD8 T cells (37)(Fig. 3D). Genetic suppression of mature T cells significantly reduced survival of KMA mice by at least the same extent as transient CD8 T cell depletion (Fig. 3E). Curiously, the opposite effect was found in KM mice, wherein deletion of *Tcrb* led to a significant extension of survival post tumour initiation (Fig. 3F). We speculate that this may reflect a pro-tumour role of a subset of T cells in the absence of an active canonical CD8 T cell response (38). Overall, these results strongly support the conclusion that the extended survival of KMA mice derives from the anti-tumour activity of tumour-infiltrating CD8 T cells.

**Figure 3:**
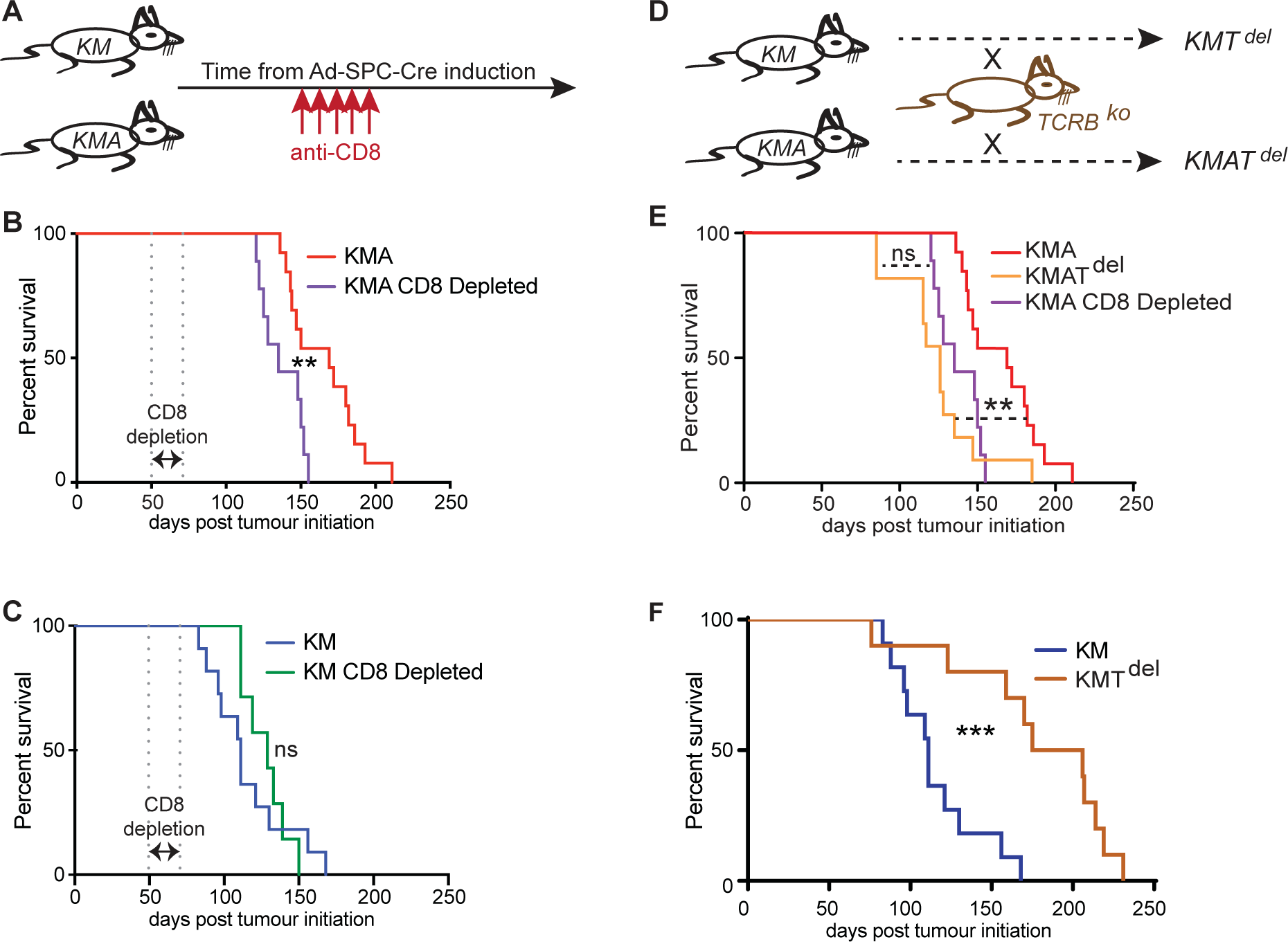
Loss of CD8+ T cells abolishes survival benefit of KMA mice. **A)** Schematic of antibody-mediated depletion of CD8+ T cells in KM and KMA mice. **B)** Overall survival of KMA mice treated with CD8 T cell depleting antibody (N= 9) compared with untreated KMA mice (N = 13, from Fig 1B). **C)** Overall survival of KM mice treated with CD8 T cell depleting antibody (N = 7) compared with untreated KM mice (N = 11, from Fig 1B). Mantel-Cox log rank test. **D)** Schematic of breeding scheme to generate KM and KMA mice with genetic deletion of *Tcrb*+ T cells (T^del^). **E)** Comparison of overall survival of KMAT^del^ (N = 12) with that of untreated KMA mice (N = 13, from Fig 1B) and CD8-depleted KMA mice (N = 9, from Fig 3B). Kruskal Wallis test. **F)** Overall survival of KMT^del^ mice (N = 10) compared with KM mice (N = 11, from Fig 1B). Mantel-Cox log rank test. For all panels, ** denotes P < 0.01; *** denotes P < 0.001.

### Neither PD1 blockade nor increased Type 1 IFN extend the anti-tumour CD8 response in KMA mice

Ligation of PD1 by PD-L1 inactivates T cells and is important for restoration of homeostasis following an acute immune response (39). Accordingly, *CD274* (encoding PD-L1) expression is induced by IFNγ produced by activated T and NK cells (40). PD-L1 is moreover implicated as a key mediator of MYC-induced suppression of anti-tumour immunity (41). Examination of *Cd274* mRNA by ISH showed strong expression at 8 weeks p.i. and somewhat lower expression at end point in both KM and KMA tumours, consistent with the presence of the MYC expressing transgene in both genotypes (Fig. 4A, B). Prolonged treatment with PD1 blocking antibody however had no impact on survival of either KM or KMA mice (Fig. 4C, D), nor did treatment result in sustained CD8 T cell tumour infiltration, examined at 12 weeks p.i. (not shown). Note that the same antibody can dramatically enhance survival in a mouse model of sarcomatoid Mesothelioma (Farahmand & Murphy, unpublished).

**Figure 4:**
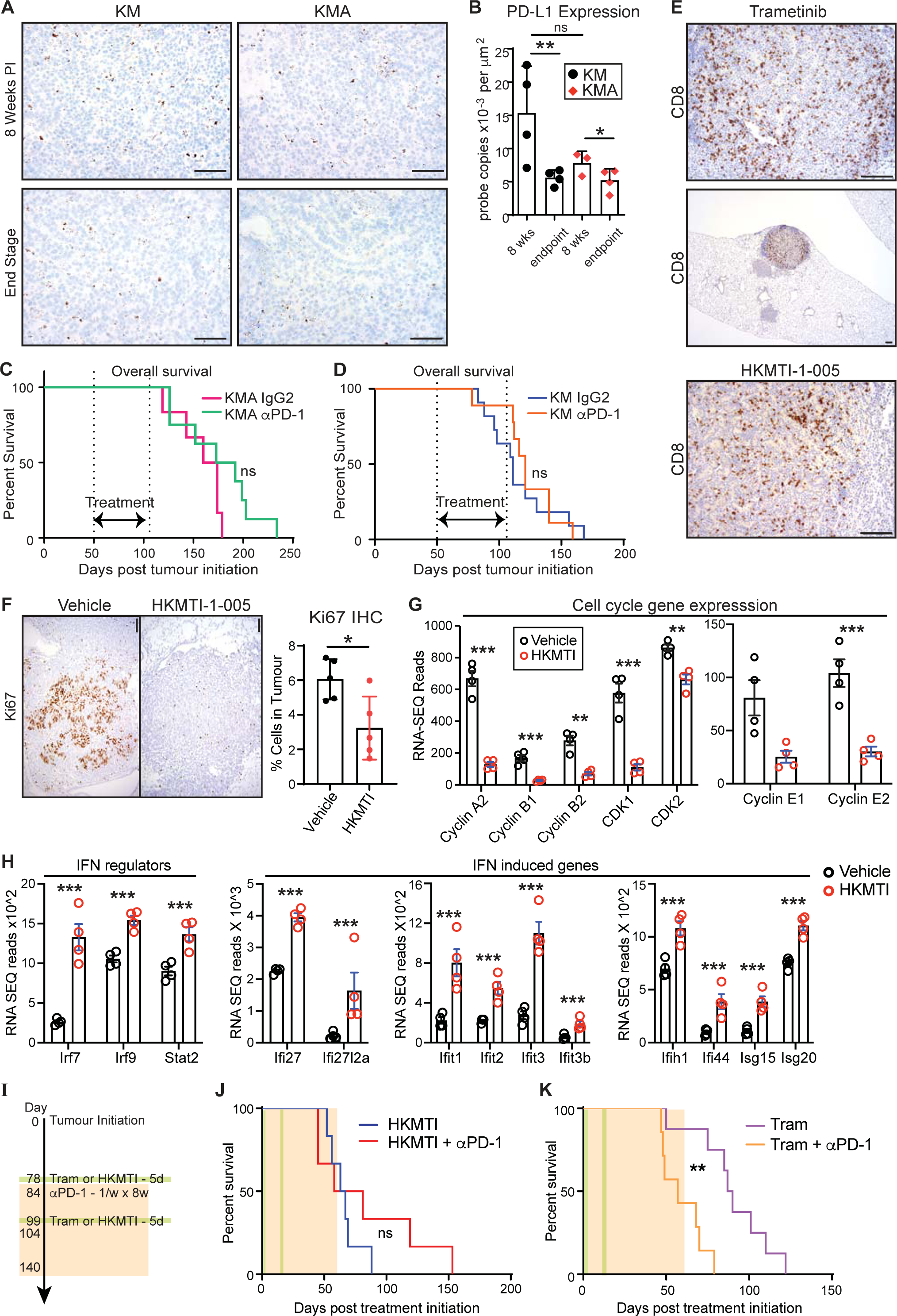
PD1 blockade and Type I IFN induction fail to extend the immune response to KMA tumours. **A)** Lung sections from KM and KMA mice at 8-weeks post allele induction and end-stage, stained via ISH for expression of *Cd274* (encoding PD-L1). Scale bar, 100μm. **B)** HALO quantification of PD-L1 expression in KM and KMA tumours from mice at 8-weeks post induction and end stage (N = 3 or 4 mice per cohort, as shown). Error bars represent Mean ± SD (ANOVA with post-hoc Tukey test). **C)** Overall survival of KMA mice treated with αPD-1 (N = 8) versus IgG2 isotype control antibody (N = 6). Mantel-Cox log rank test. **D)** Overall survival of KM mice treated with αPD-1 (N = 9) versus IgG2 isotype control antibody (N = 11, from fig 1B). Mantel-Cox log rank test. **E)** CD8α staining of super-responder KMA tumours following treatment with 1mg/kg Trametinib daily or 40mg/kg HKMTI-1-005 twice daily for 5 days prior to cull at 12-weeks post allele induction (day 78 to day 83). Scale bar, 100μm. **F)** Representative images of Ki67 stained tissue sections harvested from KMA mice treated with vehicle or 40mg/kg HKMTI-1-005 for 5 days prior to harvest at 12-weeks post allele induction. Scale bars, 100µm. HALO quantification of percentage of Ki67+ cells in lung tumours from KMA mice treated with Vehicle (N = 5) or 40mg/kg HKMTI-1-005 (N = 5) for 5 days prior to cull. Each data point represents the average score of all tumours from a section in an individual mouse. Error bars represent Mean ± SD (unpaired t-test). **G)** Normalised RNA-Seq reads of indicated cell cycle genes in tumour bearing lungs from KMA mice treated with vehicle (N = 4) or 40mg/kg HKMTI-1-005 (N = 4) for 5 days prior to cull at 12-weeks post allele induction. Mean and SEM shown. P values adjusted for multiple comparisons. **H)** Normalised RNA-Seq reads of IFN regulator and IFN-induced genes from KMA mice as per (G). **I)** Schematic of treatment schedule for KMA mice with combination treatments of Trametinib (1mg/kg) ± αPD-1 (200µg) or HKMTI-1-005 (40mg/kg) ± αPD-1 (200µg). **J)** Survival post treatment initiation of KMA mice treated with HKMTI-1-005 alone (N = 6) or HKMTI-1-005 in combination with αPD-1 (N = 6). **K)** Survival post treatment initiation of KMA mice treated with Trametinib alone (N = 8) or Trametinib in combination with αPD-1 (N = 7). Mantel-Cox log rank test. For all panels, ns denotes not significant; * denotes P <0.05; ** denotes P <0.01; *** denotes P < 0.001.

Our previous work in mouse models of pancreatic cancer revealed evasion of immunosurveillance through the combined activity of KRAS and MYC via transcriptional suppression of the immunostimulatory Type I IFN pathway, a finding since extended to several other cancer types including LuAd (42–44). MYC forms repressive transcriptional complexes with MIZ1 (encoded by *Zbtb17*) and the G9A histone methyltransferase (encoded by *Ehmt2*), while KRAS promotes MYC protein stability (45–48). We therefore asked if acute inhibition of this signalling cascade might restore CD8 T cell infiltration at the 12-week timepoint. Accordingly, KMA mice were treated for 5 days with either the MEK inhibitor Trametinib, or with an experimental G9A inhibitor, HKMTI-1-005, recently shown to stimulate Type I IFN gene expression and anti-tumour immune activity in ovarian cancer (49), and harvested on day 84. Neither drug restored CD8 T cell infiltration of the vast majority of KMA tumours examined at 12 weeks p.i., although a small number of tumours in a subset of drug-treated mice did show dramatic levels of both CD8 and CD4 T cell infiltration (Fig. 4E, S6A, B). HKMTI-1-005, however, consistently reduced cell proliferation across all lung tumours of treated mice (Fig. 4F), consistent with the established role of MYC-mediated repression in regulating the cell cycle (50). Accordingly, RNA-SEQ analysis following acute HKMTI-1-005 treatment showed a sharp reduction in cell-cycle related gene expression and a pronounced increase in Type I IFN pathway gene expression (Fig. 4G, H, S6C, D). The latter included upregulation of key IFN signal transducers *Irf7, Irf9* and *Stat2*, all direct targets MYC/MIZ1 transcriptional repression (44, 51), along with numerous IFN-responsive genes. These changes were accompanied by an apparently oligoclonal mature B cell response, evidenced by increased expression of *IghG*, indicative of immunoglobulin class switching and B cell maturation, along with multiple *IgV* genes (Fig. S6E, F). Attempts to prime an immunotherapy response by combining 2 rounds of acute treatment with either Trametinib or HKMTI-1-005 followed by α-PD1 treatment did not however extend survival of KMA mice (Fig. 4I-K). We conclude from these results that immune evasion mechanisms beyond PD-L1 expression and Type I IFN suppression are dominant in the KMA LuAd model.

### Enrichment of an EGFR/ERBB ligand-driven tumour progression signature following PD1 blockade

To investigate the failure of PD1 blockade to extend the anti-tumour immune response of KMA mice, we used single cell RNA-SEQ to examine the transcriptional response to α-PD1 in KMA tumours. Cohorts of KMA mice were treated as above with α-PD1 or left untreated, commencing at 50 days p.i., and harvested at 56 (ie. 8 wks) or 84 (12 wks) days p.i. for single-cell isolation and library generation using the 10X Genomics platform. Lung fragments containing tumours were briefly digested and separated by FACS to recover all viable cells present. Viable cells separated into 22 clusters and the ImmGen databrowser was used to establish cell identity for each cluster (Fig. 5A and Table S1). Tumour cell clusters were identified based on expression of the conditional IRFP allele, with enrichment of tumour cells from the 12-week α-PD1-treated cohort in cluster 3 and of the 12-week untreated cohort in cluster 12 (Fig. 5B, C). PANTHER overrepresentation analysis was used to directly compare clusters 3 and 12, revealing significant enrichment of “susceptibility to T cell mediated cytotoxicity” (89.4x enrichment; 2.85e^-2^ FDR) and “susceptibility to natural killer cell mediated cytotoxicity” (53.6x enrichment; 4.44e^-3^ FDR) pathways along with “EGFR/ERBB signalling pathway” (33.5x enrichment; 1.06e^-2^ FDR), following PD1 blockade (Table S2). Interrogation at the individual gene level revealed sharply higher expression of EGFR/ERBB ligands *Areg, Hbegf* and, to a lesser extent, *Ereg*, along with loss of pneumocyte differentiation markers *S*ftpa1*, Sftpb* and *Sftpc*, in 12-week mice treated with α-PD1 (Fig. 5D-F). These were accompanied by increased expression of the YAP effector *Tead1* and the Hippo pathway target *Nt5e* (encoding CD73); increased expression of inflammatory chemokines *Cxcl1, Cxcl2 , Cxcl3, Cxcl5*, and *IL24*; and reduced expression of interferon pathway and MHC II antigen presentation genes *H2-Aa*, *H2-Ab1* and *CD74* (Fig. 5E-H). Notably, increased expression of ERBB ligands and loss of lineage identity in LuAd, evidenced here by the sharp loss of pneumocyte differentiation marker expression, were previously linked to tumour progression (30, 52), and occur in a large subset of both KRAS wild-type and KRAS mutant LuAd (30, 53). Collectively, these data suggest that ERBB-mediated tumour progression may facilitate LuAd escape from immunosurveillance and is enriched by immune checkpoint blockade.

**Figure 5:**
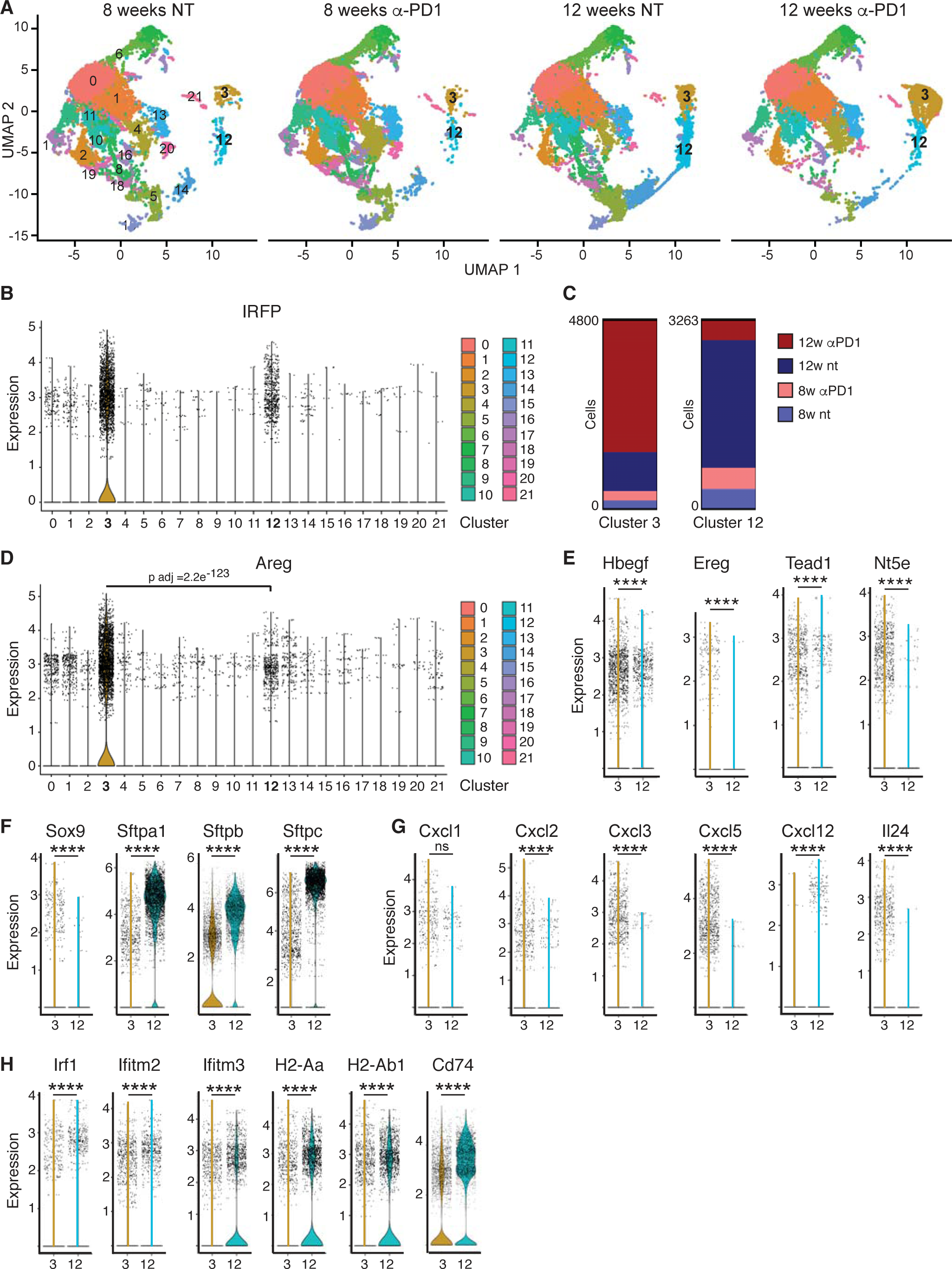
Single Cell Sequencing Analysis reveals an Enrichment of an EGFR/ERBB ligand-driven tumour progression signature following PD1 blockade. **A)** UMAPs of 22 clusters identified from single-cell RNA-Seq analysis of tumour-bearing lungs from KMA mice treated with (N=4) or without (N=4) αPD-1 and harvested at 8 or 12 weeks, as indicated. All data were pooled prior to determination of cell clusters. Clusters 3 and 12 are indicated in samples from 12-weeks untreated and 8- and 12-week αPD-1 treated samples. **B)** Violin plot of iRFP expression across all clusters identified in (A). iRFP expression identifies clusters 3 and 12 as tumour cells. **C)** Proportions of each treatment group present in clusters 3 and 12 illustrates enrichment of 12-week αPD-1 samples in cluster 3 and an enrichment of 12-week untreated samples in cluster 12. **D)** Violin plot of relative *Areg* expression across all clusters reveals a significant enrichment of Areg expression in cluster 3. **E)** Relative expression of *Hbegf, Ereg, Tead1*, and *Nt5e* in clusters 3 & 12. **F)** Relative expression of pneumocyte differentiation markers *S*ftpa1*, Sftpb*, and *Sftpc* along with the progenitor cell transcription factor *Sox9*, in clusters 3 & 12. **G)** Relative expression of inflammatory chemokines in clusters 3 & 12. **H)** Relative expression of interferon pathway and MHC II antigen presentation genes in clusters 3 & 12. (E-H) **** denotes P<0.0001; ns = not significant (T-test).

### Expression of AREG correlates with invasion in human LuAd

To investigate the relevance of these observations for human LuAd, we used multiplex immunofluorescence to examine an extensive lung tumour microarray, comprising 2653 biopsy needle cores from 1008 tumours surgically resected from treatment-naïve LuAd patients. TMAs were stained for expression of the EGFR/ERBB ligand AREG, along with Tyr^1248^-phosphorylated (ie. activated) ERBB2, and pan-Cytokeratin as a marker for epithelial differentiation. AREG expression was higher in tumour epithelium than stroma and significantly correlated with p-ERBB2 in both compartments (Fig. 6A, B). Importantly, AREG expression was significantly higher in samples from invasive, relative to non-invasive (in situ) adenocarcinoma, while tumour regions with the highest expression of both AREG and p-ERBB2 often showed loss of Cytokeratin staining, suggestive of de-differentiation and loss of lineage identity (Fig. 6C, D). These data link expression of the EGFR/ERBB ligand AREG to malignant progression in human LuAd, although it should be noted that we did not find a significant link between AREG expression and overall survival in this patient cohort.

**Figure 6:**
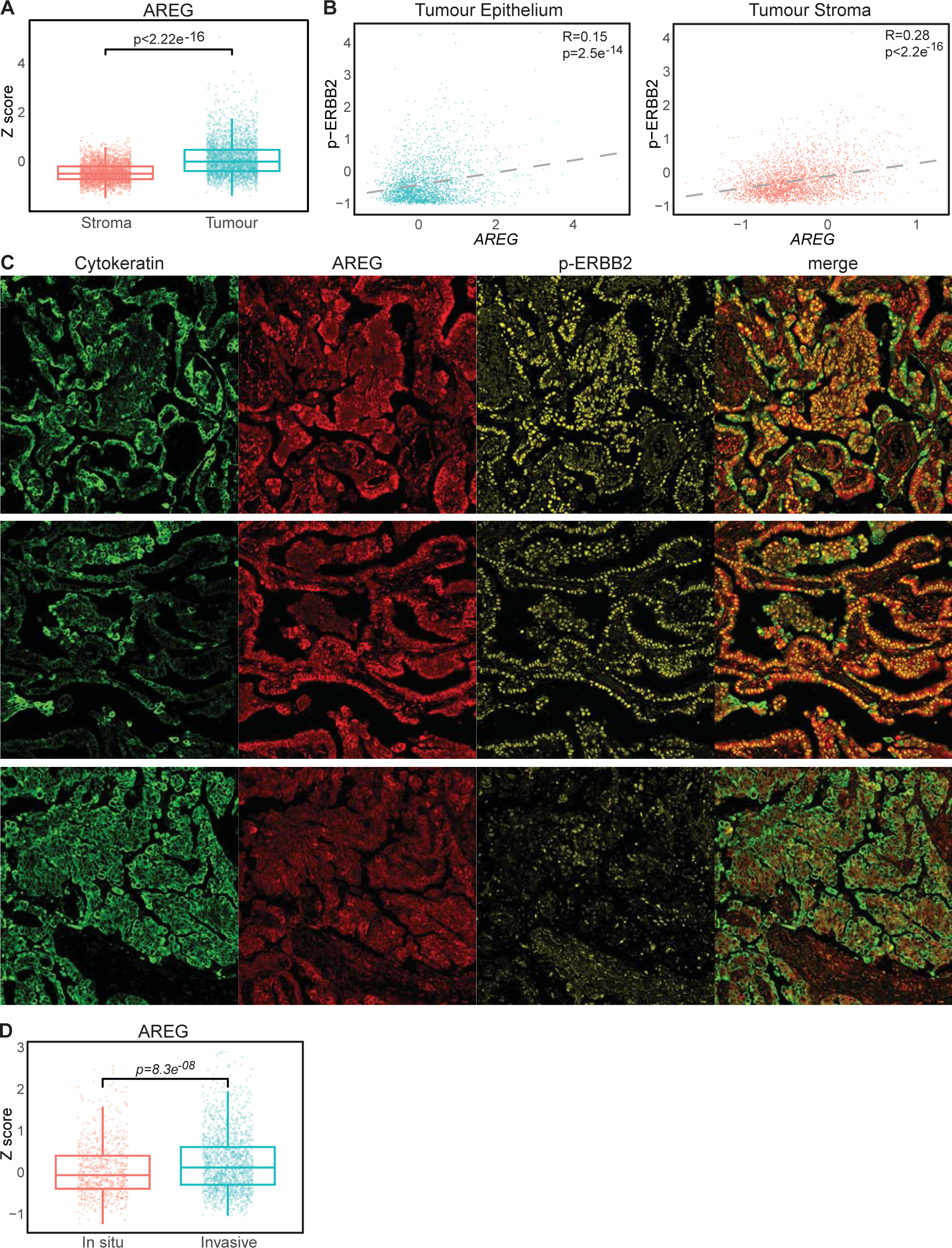
Expression of Areg correlates with invasion in human LuAd. **A)** Comparative expression of Amphiregulin (AREG) protein, segmented for tumour versus stroma, measured by IHC across a tissue microarray (TMA) of 2653 human LuAd biopsy samples (Wilcoxon Test, N=2653). **B)** Correlation of AREG protein expression and levels of phosopho-ERBB2^Y1248^ detection in LuAd tumour epithelium and associated stroma, as per (A) (Spearman test, N=2653 biopsy cores). **C)** Examples of multiplex IF images of tumour biopsies, as per (A), stained for pan-Cytokeratin, AREG and p-ERBB2^Y1248^. Top 2 rows show exemplars of high AREG/p-ERBB2 and low Cytokeratin; bottom row shows exemplar of low AREG/p-ERBB2^Y1248^, high Cytokeratin. **D)** Comparative expression of AREG as per (A) in biopsies histo-pathologically characterised as In Situ (N = 702) versus Invasive (N = 1621; Wilcoxon test).

### ERBB blockade restores anti-tumour immune activity

The ERBB family of receptor tyrosine kinases is comprised of EGFR, ERBB2, ERBB3 and ERBB4, all of which can heterodimerise with each other as well as forming homodimers. Previous work established that KRAS-driven lung tumours express *Egfr*, *Erbb2* and *Erbb3*, along with a spectrum of activating ligands including *Areg*, and that tonic signalling through the ERBB family is required for KRAS-driven lung tumour formation and increases during progression to advanced disease (30, 54). Examination of bulk RNA-SEQ data showed no significant differences in expression of *Erbb* family RTKs or ligands between KM and KMA tumours at 8 weeks post initiation (Fig. 7A). We therefore asked if ERBB blockade could reverse immune evasion by KMA tumours at 12 weeks. Accordingly, acute (5 day) treatment of KMA mice with the multi EGRF/ERBB inhibitor Afatinib resulted in widespread restoration of tumour infiltrating CD8 T cells (Fig. 7B, C) and a modest but significant reduction in Ki67 expression (Fig. 7D, E). We therefore asked if short-term Afatinib treatment might combine with PD1 blockade to enhance survival. KMA mice were treated with 2 cycles of 5 days Afatinib treatment, separated by a 2-week interval, concurrent with weekly treatment with either α-PD1 or IgG2 isotype control for up to 8 weeks, then monitored for progression to clinical endpoint (Fig. 7F). The combination of brief Afatinib priming followed by PD1 blockade significantly extended survival over mice similarly treated with Afatinib and IgG2 control, demonstrating that ERBB blockade can enhance ICB therapy *in vivo* and suggesting a plausible route to similarly enhancing ICB treatment of human LuAd.

**Figure 7:**
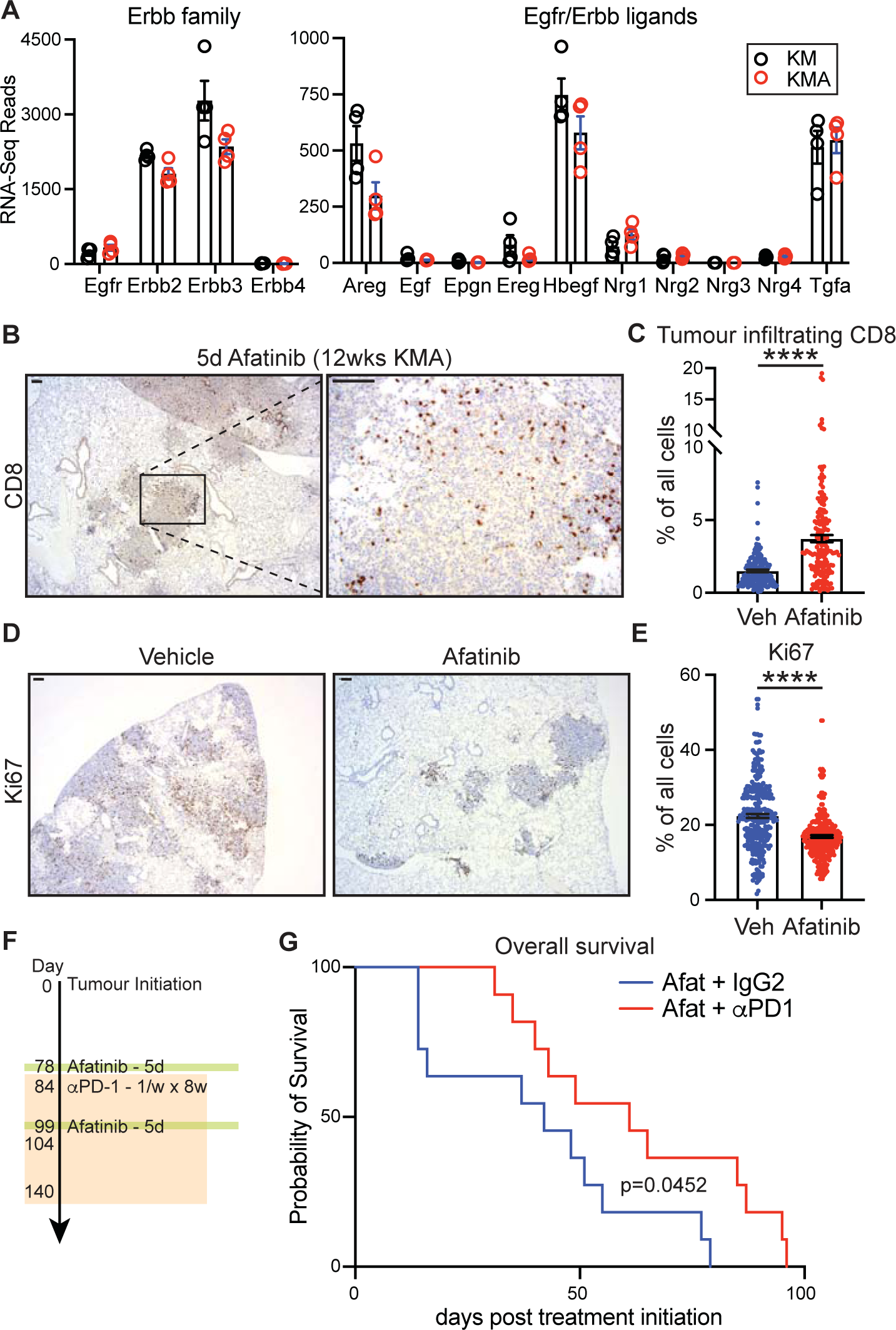
ERBB blockade restores anti-tumour immune activity. **A)** Normalised RNA-Seq reads of Erbb family genes and Egfr/Erbb ligand genes from tumour bearing lungs of KM and KMA mice harvested at 8-weeks post allele induction (N = 4 biological replicates). **B)** Representative images of CD8α stained KMA lungs following treatment with 15mg/kg Afatinib once daily for 5 days immediately prior to harvest at 12-weeks post allele induction. Scale bars = 100µm. **C)** HALO quantification of the percentage of cells in tumours that are CD8α+ following Afatinib treatment or vehicle control as per (B). Error bars represent Mean ± SEM (unpaired T-test). **D)** Representative images of Ki67 stained KMA lungs following treatment with 15mg/kg Afatinib once daily for 5 days immediately prior to harvest at 12-weeks post allele induction. Scale bars = 100µm. **E)** HALO quantification of the percentage of cells in tumours that are Ki67+ following Afatinib treatment or vehicle control as in (B). Error bars represent Mean ± SEM (unpaired T-test). **F)** Schematic of treatment schedule for KMA mice with combination treatment of Afatinib (15mg/kg) ± αPD-1 (200µg). **G)** Survival post treatment initiation of KMA mice treated with Afatinib (N = 11) or Afatinib in combination with αPD-1 (N = 11). Mantel-Cox log rank test. For all panels, * denotes P <0.05; **** denotes P < 0.0001.

## DISCUSSION

Immune checkpoint blockade (ICB) is replacing conventional therapies as the treatment of choice for lung cancer, including KRAS-mutant lung adenocarcinoma, wherein a recent study evidenced superiority of ICB over chemotherapy in real-world patients (55). Nevertheless, individual responses to ICB remain unpredictable. Co-mutations certainly contribute to response variation, with functional loss of Trp53 predicting better response while *STK11* mutation predicts resistance of lung cancers to ICB therapy (56). However, a multitude of additional factors, some known, likely many unknown, contribute further to response variation (7). A much deeper mechanistic understanding of the dynamics of tumour and anti-tumour immune co-evolution is required to fully exploit the potential of cancer immunotherapy. Heretofore, a lack of suitable model systems in which to interrogate these dynamics in an autochthonous setting has hampered progress. The development in recent years of immunogenic mouse models of sporadic lung cancer (eg. (14, 17, 26, 57) has begun to address this limitation, but no individual model can possibly reflect the spectrum of human disease heterogeneity, and multiple models will be needed to tile across the disease-positioning space for any given cancer. In this regard, we report here the development of three new mouse models of progressive lung adenocarcinoma, KMA, KPA and KLA, and show that co-expression of APOBEC3B alongside KRAS^G12D^ and MYC drives a transiently effective CD8 T cell-mediated immune response leading to a significant extension of lifespan over comparator mice lacking the *Rosa26^DS.lsl-APOBEC3B^* (A3B) allele. Mirroring the varied sensitivities of human KRAS-driven LuAd to ICB (56), a similar extension of survival was measured when A3B was combined with *p53* loss in KPA mice, but not when combined with loss of *Stk11*, encoding Lkb1, in KLA mice. A lack of sufficient tumour material at 8 weeks post induction precluded investigation of the immune response in KPA mice. It is therefore curious that tumour burden in end-stage KPA mice closely resembled that of KP comparators which succumb much earlier, suggesting that tumour initiating cells in the KPA model somehow evade immune clearance and emerge following the initial immune response to give rise *de novo* to tumours, long after Cre-mediated allelic recombination. Further work on the KPA model will shed light on mechanisms of such apparent tumour dormancy and evasion of immune clearance by tumour initiating cells, whereas the KLA model will provide a useful platform for investigating intrinsic resistance to anti-tumour immunity.

We expected APOBEC3B to drive immune visibility through a mutagenic process arising from the enzyme’s Cytidine deaminase activity combined with inaccurate DNA repair mechanisms (24). Although we verified increased Cytidine deaminase activity in cell lines generated from end-stage KPA tumours, efforts to generate KMA cell lines were unsuccessful and we were unable to measure consistently increased mutation burden in primary end-stage tumours from KMA mice. A separate study, using an independently generated *Rosa26-lsl-A3B* allele with low expression of A3B similar to ours, also found no evidence of increased mutagenesis in primary tumours but did identify increased mutation burden and evidence of Cytidine mutations in cell lines derived from end-stage KPA tumours (26). A preprint available at the time of writing has now shown that much higher levels of A3B expression, driven from the artificial CAG promoter-enhanced *Rosa26* locus, suffice to generate tumours in mice after a protracted latency period of up to 600 days (35). Importantly, this latter study reports successful identification of the canonical C->T APOBEC mutagenesis signature, predominantly in the context of TC dinucleotides conforming to COSMIC base substitution signature SBS02 (36), in tumours from *Rosa26.CAG-A3B* mice. Together, this would suggest that A3B mutagenesis only measurably manifests upon clonal outgrowth of cells wherein such mutations generate a survival advantage and, given the much shorter timeframe of our study using strong oncogenic driver mutations, the level of mutagenesis present when A3B is driven from the unenhanced (ie. endogenous) *Rosa26* locus fails *in vivo* to confer any clonal advantage that would allow detection of such signatures. While we cannot formally rule out the possibility that our T cell mediated response is directed against the human APOBEC3B protein itself, we note that end-stage KMA tumours continue to express the *APOBEC3B* transgene, as do cell lines derived from end-stage KPA tumours, arguing against immune editing of tumour cells expressing A3B *per se*. Moreover, reactivation of endogenous retroviruses was recently shown to drive immunogenicity in a similar model of A3B-expressing mutant KRAS-driven lung cancer, although the precise mechanism of A3B-induced retroviral reactivation remains obscure (58).

KMA tumour infiltration by both CD4 and CD8 T cells was prominent at 8 weeks post tumour initiation but almost entirely absent by 12 weeks, yielding a unique opportunity to investigate this important dynamic *in situ*. Given strong expression of *Cd274* (encoding PD-L1) throughout the lung tumours, we hypothesized that PD1 blockade might sustain tumour infiltration if administered during this period, but this was not the case and no impact of single-agent PD1 blockade on overall survival was measured. We wondered if suppression of Type I Interferon signalling by the combined effects of KRAS and MYC might explain the lack of impact of PD1 blockade (44, 51). Acute treatment of mice with the MEK inhibitor Trametinib restored T cell infiltration in a rare subset of KMA tumours but the vast majority of tumours showed no such infiltration and, somewhat surprisingly, RNA-SEQ did not reveal the expected increase in Type I Interferon pathway activity. This contrasts with the effect of direct inhibition of KRAS^G12C^ in a transplantable cell line model of LuAd (26, 44). More surprisingly, whereas Trametinib extended survival of KMA mice when administered alone, the inclusion of PD1 blockade completely negated the survival benefit of Trametinib. The reason for this negative interaction is presently unclear and may relate to increased ERBB signalling following PD1 blockade, but the data serve to caution strongly against the use of such a combination in human patients.

We next investigated use of HKMTI-1-005 (59), an experimental dual inhibitor of G9A, which forms part of the MYC repressor complex (46), and EZH2, suppression of which was recently shown to destabilise MYC protein (60). As found with Trametinib treatment, a rare subset of KMA tumours showed pronounced CD8 T cell infiltration following acute treatment with HKMTI-1-005. Moreover, treatment of KMA mice with HKMTI-1-005 strongly inhibited tumour cell proliferation, and RNA-SEQ analysis revealed a clear increase in Type I IFN pathway expression along with a pronounced reduction of cell-cycle related genes, suggesting potent suppression of MYC activity. Mindful of the likely adverse effects of prolonged systemic MYC suppression, we again attempted to prime an immune response by treating briefly with HKMTI-1-005 followed by sustained PD1 blockade, but again failed to observe any benefit of the combination over brief treatment with HKMTI-1-005 alone, despite activation of the Type I IFN pathway by HKMTI-1-005. The absence of a T cell mediated response despite IFN restoration is perhaps not surprising, given that our previous work in pancreatic cancer showed a B cell and NK cell-dependent extension of survival, but no increase in tumour-infiltrating T cells, following restoration of IFN signalling by genetic disruption of the MYC/Miz1 repressor complex (51). Additionally, MYC is required for T cell proliferation following TCR engagement (61) and it is possible that HKMTI-1-005 treatment impedes this response.

We then used single-cell RNA-SEQ to investigate the apparent resistance of KMA tumours to PD1 blockade. Importantly, this analysis was performed on all viable cells without pre-selection of specific cell types following lung fragment digestion. Although tumour cells were somewhat underrepresented due to loss of viability during sample processing, we identified 2 clusters of tumour cells based on expression of the conditional *IRFP* and human *MYC* alleles, one (cluster 3) enriched for cells from α-PD1-treated mice harvested at 12 weeks, the other (cluster 12) enriched for cells from untreated mice harvested at the same timepoint. Focussing on these two clusters, among the topmost differentially enriched pathways, “susceptibility to T cell mediated cytotoxicity” and “susceptibility to NK cell mediated cytotoxicity” suggest a selective pressure to evade immune-mediated killing following PD1 blockade. Surprisingly, these signatures were accompanied by significant enrichment of “ERBB2-EGFR signalling pathway”, “positive regulation of epidermal growth factor-activated receptor activity” and “ERBB2 signalling pathway”. We and others previously showed that spontaneous progression of KRAS^G12D^-driven tumours to locally invasive disease in mice involves increased expression of multiple EGFR/ERBB ligands, and that signalling through endogenously expressed wild-type EGFR/ERBB is required for development of mutant KRAS-driven tumours (30, 54). Corroborating our human TMA data, ERBB ligands, including *AREG*, are highly expressed in treatment-naīve human LuAd, reflecting an alternative means of RAS pathway upregulation and associated with suppression of lineage identity (52, 53). Interestingly, the 50-gene tumour progression signature we previously identified includes multiple known immune modulators, including *Nt5e* (encoding CD73), *Ptges* (encoding Prostaglandin E Synthase), *Tnfrsf12a* (encoding the Tweak receptor) along with multiple *S100* genes (30, 62). *NT5E* encodes a cell surface enzyme that converts extracellular AMP to free Adenosine, a potent suppressor of T cell and NK cell cytotoxicity (63), and expression of *NT5E* is sharply suppressed upon treatment of human LuAd cell lines with the EGFR/ERBB inhibitor, Neratinib (30). Collectively, these results suggest that increased EGFR/ERBB signalling, driven by elevated expression of multiple EGFR/ERBB ligands, may actively participate in immune evasion. Accordingly, acute treatment with the EGFR/ERBB inhibitor Afatinib broadly restored infiltration of KMA tumours by CD8 and CD4 T cells. The subsequent addition of α-PD1 yielded a significant extension of survival over brief treatment with Afatinib alone, the first time in our hands that PD1 blockade showed any survival benefit in the KMA model.

Clinical trials of EGFR inhibitors combined with ICB in lung cancer have to date yielded rather disappointing results, however such trials have largely been limited to EGFR-mutant LuAd and focussed on use of highly EGFR-selective agents, such as Gefitinib and Osimertinib (64). Given that α-PD1 resistance in our model is associated with elevated expression of EGFR/ERBB ligands, which may engage EGFR heterodimers with ERBB2 or ERBB3, or ERBB2/3 heterodimers at high localised ligand concentrations, broad spectrum inhibition of the ERBB family would be more effective than selective targeting of EGFR alone, as we previously showed for development of KRAS^G12D^-driven lung tumours (30). Moreover, our data were generated using 2 brief rounds of Afatinib treatment to trigger α-PD1 responsiveness, suggesting that brief transient pulses of ERBB blockade can combine well with ICB and may thereby mitigate some of the clinical toxicity associated with prolonged EGFR inhibition in the context of ICB therapy (64). Further refinement of dosing schedules may thus yield greater therapeutic benefit.

In summary, we have generated a series of GE mouse models that will facilitate mechanistic investigation into multiple aspects of the dynamic co-evolution *in situ* of nascent lung tumours and their corresponding immune interactions, including intrinsic and therapy-induced resistance, and evasion of immune surveillance through dormancy. Our detailed investigation of the KMA model reveals active evasion of anti-tumour immune responses through upregulation of ERBB ligands and demonstrates the potential for broad spectrum ERBB inhibition to re-engage immune activity in ICB-refractory KRAS mutant lung tumours.

## MATERIALS & METHODS

### Mice and *In Vivo* Procedures

The *LSL-KRAS^G12D^* (65), *Rosa26^DM.lsl-MYC^* (30), *Hprt-lsl-IRFP* (*66*)*, Trp53^loxP^* (67), *Stk11^tm1.1Sjm^* (28), and B6.129P2-*Tcrb^tm1Mom/J^*(37) alleles were previously described. Generation of *Rosa26^DS.lsl-APOBEC3B^* is described in the Supplementary Material. All mice were maintained on mixed FVBN/C57Bl6 background, housed on a 12-hour light/dark cycle, and fed and watered ad libitum. Male and female mice were included in all analyses in approximately equal numbers. Recombinant adenovirus expressing Ad5mSPC-CRE (AS-CRE) was purchased from the University of Iowa gene therapy facility. To initiate lung tumours, 8- to 10-week-old mice were sedated with a mixture of medetomidine and ketamine, injected IP, followed by intranasal administration of AS-CRE. For most experiments, 1x10^8^ viral pfu AS-CRE were administered intranasally using the calcium phosphate precipitation method, as described previously (29). For lower tumour burden, 1x10^7^ pfu were administered. Mice were aged to early or intermediate timepoints in tumour development. For overall survival analysis, cohorts of mice were monitored by facility personnel with no knowledge of experimental design and euthanized when humane endpoint was reached. All mice were euthanised humanely by CO_2_ inhalation followed by cervical dislocation. For histologic analysis, mouse lungs were perfusion-fixed in 10% neutral buffered formalin overnight. Prior to fixation, a small portion of tumour bearing lung (approximately 30ug) was snap-frozen for RNA or western blot analysis from a subset of mice. To determine the incidence of metastasis to the liver, the dissected livers of end stage mice induced at 1x10^7^ pfu were visualised by PEARL infrared imaging, as previously described (66). For all intervention analyses, mice were randomly assigned to treatment or control groups, balanced only for animal sex. To determine the influence of CD8+ T cells, cohorts of randomly selected KM and KMA mice were treated with anti-mouse CD8α (200µg, i.p., *InVivo*MAb, Clone 2.43) from 50 days post allele induction, twice weekly for 3 weeks, and euthanised at clinical endpoint. To verify CD8α T cell depletion following the antibody treatment, blood was sampled by tail bleed from mice prior to and after 2 weeks of treatment. Cells were stained with CD3 (100203, Bioloegend), CD4 (100451 Biolegend), CD8 (563786, Biolegend), CD19 (47-0193-85, eBioscience) and Zombie NIR10311 (423106, Biolegend) after red blood cell lysis and analysed by flow cytometry (BD Fortessa). FACS profiles were generated and quantified using FlowJo (Tree Star). To assess the potential survival benefit of immune checkpoint blockade, KM and KMA mice were treated with anti-mouse PD1 (200µg, i.p., *InVivo*MAb, CD270, Clone RMP1-14) or isotype control (200µg, i.p., Rat IgG2α, *InVivo*MAb Clone 2A3) from 50- or 84-days post allele induction, once weekly for 8 weeks, and euthanised at clinical endpoint. To assess effects of short term PD1 blockade, mice were treated from 50 days post allele induction, once weekly for 5 weeks and euthanised at 12 weeks post allele induction. To assess efficacy of PD1 blockade in KMA mice by single cell RNA sequencing analysis, mice were treated daily with anti-mouse PD1 or IgG2 isotype control for 3 consecutive days prior to 8- and 12-weeks post allele induction respectively and euthanised. To assess the acute impact of targeting the MYC/G9A transcriptional repressor complex in KMA mice, HKMTI-1-005 (40mg/kg) or vehicle control (0.9% NaCl) were administered by twice daily i.p. injection for 5 consecutive days prior to 12 weeks post allele induction and euthanised. Potential survival benefit of HKMTI-1-005 was assessed in KMA mice with and without PD1 blocking antibody. Mice were treated with two 5-day pulses of 40mg/kg HKMTI-1-005, starting at days 79 and 100 post allele induction, and 8 weeks of once weekly αPD1 from day 84 post allele induction and euthanised at clinical endpoint. To assess the acute impact of MEK inhibition in KMA mice, Trametinib (1mg/kg) (Medchem Express) was administered by single daily i.p. injection for 5 consecutive days prior to 12 weeks post allele induction and euthanised. Potential survival benefit of Trametinib was assessed in KMA mice with and without αPD1. Mice were treated with two 5-day pulses of 1mg/kg Trametinib at days 79 and 100 post allele induction, and 8 weeks of once weekly αPD1 as described above from day 84 post allele induction and euthanised at clinical endpoint. To assess the impact of acute multi-ERBB inhibition in KMA mice, Afatinib (15mg/kg) (Medchem Express) or vehicle control (0.5% hydroxypropyl methycellulose and 0.1% Tween80 in H_2_O) was administered by single daily oral gavage for 5 consecutive days prior to 12 weeks post allele induction and euthanised. Potential survival benefit of Afatinib was assessed in KMA mice with and without PD1 blockade. Mice were treated with two 5-day pulses of 15mg/kg Afatinib at days 79 and 100 post allele induction, and 8 weeks of once weekly αPD1 as described above from day 84 post allele induction and euthanised at clinical endpoint.

### Immunohistochemistry and Tissue Analysis

All mouse tissue IHC and ISH staining was performed on 4µm formalin-fixed, paraffin-embedded sections, which had been previously heated to 60°C for 2 hours. Peroxidase blocking was performed for 10 minutes in 1% H_2_O_2_ diluted in H_2_O, followed by heat-mediated or enzyme-mediated antigen retrieval. Nonspecific antibody binding was blocked with up to 3% BSA or 2.5% normal horse serum for one hour at room temperature. The following antibodies were used at the indicated dilution and indicated antigen retrieval method: Ki67 (CST 12202, 1:1000, ER2 Leica), CD8a (eBioscience 14-0808-82, 1:500, ER2 Leica), CD4 (eBioscience 14-9766-82, 1:500, ER2 Leica), FoxP3 (CST 12653, 1:200, pH6), Nkp46 (R & D Systems aF2225, 1:200, pH6), F4/80 (Abcam ab6640, 1:100, Enzyme 1 Leica), CD45R (Abcam ab64100, 1:200, ER2 Lecia), and Ly6G (2B Scientific BE0075-1, 1:60000, ER2 Leica). Nkp46 and FoxP3 were stained on a Dako AutostainerLink48. CD8a, CD4, CD45R, F4/80, Ly6G and mouse IgG were stained on the Leica Bond RX Autostainer. Rabbit Envision (Agilent), Goat ImmPRESS and Rat ImmPRESS kits were used as secondary antibodies. Mouse envision (K4001, Agilent) diluted 1:3 with PBS was used for detection of IgG. Horseradish peroxidase (HRP) signal was detected using liquid DAB (Agilent or Leica). Sections were counterstained with hematoxylin and cover-clipped using DPX mount (CellPath). ISH detection of *Hs-Apobec3b* and mm-*CD274* (Advanced Cell Diagnostics) mRNA was performed using RNAscope 2.5 LS Detection Kit (ACD) on a Leica Bond Autostainer, strictly adhering to ACD protocols. Tumour area was calculated using HALO software (Indica Labs) as the percent area of the lung occupied by adenocarcinoma, measured on hematoxylin and eosin stained sections. Immune infiltration was quantified using HALO software in the case of F4/80, Ly6G and FoxP3. Immune infiltration was quantified using QuPath software in the case of CD8, CD4 and NK cells.

### Bulk Tumour RNA Sequencing

RNA was isolated from flash frozen lung tumour samples using Qiagen RNeasy Kit according to the manufacturer’s protocol for purification of total RNA from animal tissues. DNA was depleted with the RNase-free DNase Set (Qiagen). The quality of purified RNA was tested on an Aligent 2200 Tapestation using RNA screentape. Libraries for cluster generation and DNA sequencing were prepared following an adapted method from (68), using Illumina TruSeq Stranded mRNA LT Kit. Quality and quantity of the DNA libraries were assessed on an Aligent 2200 Tapestation (D1000 screentape) and Qubit (Thermo Fisher Scientific) respectively. The libraries were run on the Illumina Next Seq 500 using the High Output 75 cycles kit (2 x 36 cycles, paired end reads, single index).

### Single Cell RNA Sequencing

Lungs were dissected from mice treated as described above and a tumour bearing lobe mechanically dissociated and transferred to a collagenase solution (DMEM medium (Thermo Fisher Scientific), 1mg/ml collagenase D (Roche) and 25µg/ml DNase 1 (THermo Fisher Scientific)). The gentleMACS Octo Dissociator (Miltenyi Biotech) was used to perform enzymatic dissociation, run: 37C_M_LDK_01, according to the manufacturer’s dissociation protocol. Lung suspensions were filtered through a 70-µm strainer with a syringe plunger and enzymatic reaction was stopped by addition of 5ml DMEM supplemented with 10% FCS, 2mM L-glutamine (Thermo Fisher Scientific) and 10,000 U/mol penicillin/streptomycin (Thermo Fisher Scientific). Cells were suspended in 5ml of 1x Red Blood Cell Lysis Buffer (Thermo Fisher Scientific) for 3 minutes. Cells were resuspended in PBS containing 0.5% BSA and counted using a hemocytometer. A total of 40,000 cells were loaded onto each channel of Chromium Chip G using reagents from the 10x Chromium Single-Cell 3’ v3 Bead Kit and Library (10X Genomics) according to the manufacturer’s protocol. The libraries were analysed using the Bioanalyzer High Sensitivity DNA Kit (Aligent Technologies). scRNA-Seq libraries were sequenced on the Illumina NovaSeq 6000 with paired-end 150-base reads.

### Deaminase Assay & AP Sites Quantification

KP cells (368-T1 and 394-T4) were gifted from Tyler Jacks. KPA cells were generated from tumour-bearing KPA mice in-house. Cells were grown in RPMI 1640 with 10% FCS, 2mM L-glutamine (Thermo Fisher Scientific) and 10,000 U/mol penicillin/streptomycin (Thermo Fisher Scientific). Cells were lysed in 25mM HEPES (pH7.4), 10% glycerol, 150mM NaCl, 0.5% Triton X-100, 1mM EDTA, 1mM MgCl2 and 1mM ZnCl2, supplemented with protease inhibitors. Lysates were incubated at 37°C for 15 minutes following addition of 2µg RNase A (Qiagen). 1pmol ssDNA substrate 5’ DY782 ATTATTATTATTATTATTATTTCATTTATTTATTTATTTA-3’ (Eurofins, UK) and 0.75U uracil-DNA glycosylase (NEB) were added to 10µg protein lysate at 37°C for 1 hour. 10µl 1N NaOH was added and samples incubated for 15 minutes at 37°C. Finally, 10µl 1N HCl was added to neutralise the reaction and samples were separated by electrophoresis through 15% urea-PAGE gels in TBE at 150V for 1 hour. Deamination activity was determined by densitometric quantification of cleaved substrate using ImageJ software. Data shown is from three independent experiments. AP sites were quantified using AP sites quantification kit (Cell Biolabs, Inc.) according to the manufacturers instructions. DNA from three independent samples was used for the assay.

### Immunoblotting

Denaturing PAGE was conducted using standard protocols. Primary antibodies for western blot were used at the following dilutions – MYC (Y69) (Abcam 32072, 1:1000), KRAS^G12D^ (CST 14429, 1:400), Pan-RAS (Cytoskeleton AESA02, 1:250), APOBEC3B (PA6889, gifted from Simak Ali, ICL, London 1:500), Vinculin (Santa Cruz 25336, 1:1000). Secondary horseradish peroxidase conjugated antibodies were added at a concentration of 1:7000 in 5% milk in TBST (HRP anti-mouse NA931V; HRP antirabbit NA934V, both GE Healthcare) and detected using chemiluminescence Clarity or Clarity Max ECL western blotting substrate (Biorad 1705061 or 1705062).

### Statistical Analyses

Raw data obtained from HALO or QuPath was copied into GraphPad Prism spreadsheets. All Mean, SD and SEM values of biological replicates were calculated using the calculator function. Graphical representation of such data was produced in GraphPad Prism. Statistical significance was determined by the Student T test. For multiple comparisons, ANOVA was used with a post hoc Tukey test or post hoc Fischer LSD test. For non– normally distributed data (e.g., survival benefit and immune cell infiltration), Mantel–Cox (two-way comparison), or Kruskal–Wallis (multiple comparison) tests were performed. For bulk RNA-SEQ data, adjusted P values calculated in R are shown. For single cell RNA-SEQ pathway enrichment analysis Fisher Test with false discovery rate correction was used. For all significant results shown * denotes P < 0.05; ** denotes P < 0.01; *** denotes P < 0.001, **** denotes P < 0.0001.

### Data Availability

Large data sets (with the exception of clinical and confidential data) will be made publicly available at the time of publication, if not sooner. These primary and processed data and analysis pipelines will be accompanied by appropriate metadata, in accordance with FAIR data principles and repository requirements.

## ETHICS STATEMENT

Procedures involving mice were authorised by the local animal welfare committee, AWERB, performed in accordance with Home Office Licence numbers 70/7950 & PE47BC0BF (CRUK BICR, UK), and adhered to the principles of the NC3Rs.

## Supporting information

Supplemental information incl 6 figures

## ACKNOWLEDGEMENTS

The authors thank all members of the Murphy lab for vibrant discussion of the project throughout development; all the staff of the CRUK Beatson Biological Services Unit (BSU), Histology core facility and the Beatson Advanced Imaging Resource (BAIR); Prof. Reuben Harris for provision of APOBEC-3B antibodies; CRUK Beatson Integrity Officer Catherine Winchester for critical reading of the manuscript and guidance on best publishing practice. For the purpose of open access, the author(s) has applied a Creative Commons Attribution (CC BY) licence to any Author Accepted Manuscript version arising from this submission.

## AUTHOR CONTRIBUTIONS

*Conceptualization* DJM, MRJ, DS, SBC

*Data Curation* SL, RS, RK, SM, IP

*Formal Analysis* SL, BK, RS, LOJ, SE, IP, DMK, YCH, RP, NB, RK, SM, MP, ICD, ER, DJM

*Funding Acquisition* DJM, JLQ, SBC

*Investigation* SL, BK, SE, SM, MP, ICD

*Methodology* SL, BK, LOJ, SE, DMK, YCH, RP, NB, AL, MP, ICD, CN, GC, KK

*Resources* MRJ, SA, MJF, DN, KK, RBB, JLQ, DS

*Software* n/a

*Supervision* MRJ, CM, SA, DN, KK, RBB, JLQ, DS, SBC, ER, DJM

*Validation* IP, DM, DN, SC, ER *Visualization* SL, SM, IP, MP, ER, DJM *Writing – Original draft* SL, DJM

*Writing – Reviewing & Editing* All authors

