## Supplemental information incl 6 figures for "ERBB signalling contributes to immune evasion in KRAS-driven lung adenocarcinoma"

#### SUPPLEMENTARY MATERIALS

**Supplementary Methods**

**Supplementary References**

**6 Supplementary Figures**

Figure S1: Characterisation of APOBEC3B expressing tumours

Figure S2: APOBEC3B mutagenesis & Cytidine deaminase activity

Figure S3: Immune related gene expression in KMA tumours

Figure S4: Immune landscape of KM and KMA tumours

Figure S5: Depletion of CD8 T cells negates KMA survival advantage

Figure S6: Gene expression changes induced by inhibition of the MYC repressor complex

**2 Supplementary Tables**

Table S1: Cell Lineage Assignments

Table S2: Pathway Enrichment

#### SUPPLEMENTARY METHODS

##### Generation of Rosa26DS-lsl-APOBEC3B mice

Inducible expression of APOBEC3B in mice was achieved by inserting a lox-stop-lox APOBEC3B transgene into the Gt(ROSA)26Sor locus (Ensembl ID: ENSMUSG00000086429 in Genome Assembly GRCm38.p6). The human APOBEC3B sequence was codon optimised and imported as a synthetic fragment (GeneArt, ThermoFisher). The resultant APOBEC3B coding sequence was excised and subcloned into the MCS of phprtCAG.STOP. The AscI-PacI fragment (containing the CAG-lox-STOP-lox- APOBEC3B -pA element) was then excised and cloned into pROSA26-1, comprising homology arms for the Gt(ROSA)26Sor locus (Srinivas '01). The pROSA26-1-LSL- APOBEC3B targeting vector was linearised by AscI digestion and transfected into HM1 mESCs [1]. 8x106 mid-log phase HM1 ES cells (Thermo Fisher) were resuspended in Embryomax electroporation solution (Millipore) and mixed with 40µg of AscI-linearised targeting vector. Electroporation was performed under standard conditions (250V; 500µF; infinite resistance; cuvette width: 4mm) in a Biorad GenePulser XCell with capacitance extender. Under these conditions, the time constant is close to 7 msecs. After plating onto DR4 irradiated MEF (Thermo Fisher) monolayers across four 10cm plates, cells were maintained under regular ESC medium for 40-48 hours before being placed under G418 selection (300µg/ml). Cells were selected under G418 for 5-7 days, at which time, surviving colonies were picked and screened for targeting by long range PCR from within the neomycin-resistance cassette to sequences beyond the ends of the homology arms. PCR genotyping was done using Expand Long Template (Roche) according to the manufacturer's recommendations. Oligo sequences used to screen cells to ensure appropriate targeting of the Gt(ROSA)26Sor locus were ACGACCGCAGTTCCTATGAC and CGCCTAAAGAAGAGGCTGTG for the 5 prime side and CACTGACCATCATGCCTCTG and TAGTTGCCAGCCATCTGTTG for the 3 prime side. Following identification of correctly targeted clones, mouse lines were generated by injection of ES cells into C57BL/6J blastocysts according to standard protocols (Nagy et al. 2003). After breeding of chimeras, germline offspring were identified by coat colour and the presence of the modified allele was confirmed with primers specific for the optimised Apobec3b transgene (ACTGTACAAAGAGGCCCTCC and GCAACTAGAAGGCACAGTCG; 248bp). All genotyping was subsequently performed by Transnetyx Inc.

##### RNA-Sequencing Analysis

For bulk tumour RNA-SEQ, FastQ files were generated from the sequencer output using Illumina's bcl2fastq and quality checks on the raw data were done using Fast QC and Fastq screen. Alignment of the RNA-seq paired end reads was to the GRCh38 version of the mouse genome and annotation using Hisat2. Expression levels were determined and statistically analysed by a work-flow combining HTSeq, the R environment, utilising packages from Bioconductor data analysis suite and differential gene analysis based on negative binomial distribution using the DESeq2 package. Further analysis and visualisation used R and Bioconductor packages. Pathway analysis was completed using the MetaCore Software from Clarivate Analytics.

For scRNA-Seq, quality checks and trimming on the raw scRNA-Seq fastq.gz data files were done using FastQC version 0.11.9 (1), FastP version 0.20.1(2) and FastQ Screen version 0.14 (3). The creation of the transgenic reference genome and annotation was formed from the combining of the GRCm39.103 version of the mouse genome and annotation (4), the Human gene sequences for MYC, APOBEC3B and KRAS-G12D based on GRCh38.103, and the sequence for iRFP. Alignment to the transgenic reference genome and aggregation was completed using 10x Genomics Cell Ranger version 5.0.1 (5). Quality control, integration of data, clustering, marker gene identification and exploratory analysis was accomplished using the Seurat package version 4.0.4 (6) and the R environment version 4.0.3 (7) . Identification of high-quality cells used the adaptive thresholds method as outlined in Orchestrating single-cell analysis with Bioconductor (8). Conditions were integrated and analysed to identify shared cell types in each condition. Cell types were assigned through identification of conserved cell type markers across conditions, using the ImmGen online cell identifier tool (http://rstats.immgen.org/MyGeneSet_New/index.html). Proportions of cell types constituted by each condition were determined and cell types enriched in specific conditions were analysed to identify marker genes. Two tumour cell clusters were identified through expression of iRFP and these clusters were compared for differentially expressed genes. Enrichment for GO terms was carried out using Panther (12). Computational analysis was documented at each stage using MultiQC(9), Jupyter Notebooks(11) and R Notebooks (12).

**Whole Exome Sequencing and Analysis**

For next generation sequencing, exome capture was performed using the SureSelectXT mouse exon kit (Agilent). Exome capture libraries were then sequenced on a HiSeq 2000 (Illumina) using the HiSeq sequencing Kit (200 cycles). 60 M (2 × 75 bp) exome reads were sequenced from each sample. Reads were processed and variants called using the GATK4.1.4.1/Mutect2 “tumor/normal” workflow with default settings (https://www.nature.com/articles/ng.806, genome build GRCm38), considering only “PASS” variants for downstream analysis. Mutational signature analysis was performed using the MutationalPatterns R package v3.6.0 (13) and COSMIC Signatures v3.3.1, mm10 (14). We used the fit_to_signatures function to match observed mutations to reported signatures. For visualization, only signatures contributing to more than 10% of observed mutations per sample are shown.

**Tissue Microarray & Multiplex Immunoflourescence**

A fully automated multiplex immunofluorescence (mIF) assay was applied to 23 human lung adenocarcinoma FFPE tissue microarrays (TMA) containing 3075 1mm cores corresponding to 1025 patients from the LATTICeA cohort as previously described (15). Briefly, a mIF assay was developed on the Ventana Discovery Ultra autostainer (Roche Tissue Diagnostics) using the following antibodies and corresponding dilutions; Anti-ErbB2 / HER2 (phospho Y1248) (Abcam ab101229, 1:25), CD68 (D4B9C) XP® (Cell Signaling Technologies 76437, 1:200), Cytokeratin Multi (AE1/AE3) (Leica Biosystems AE1/AE3-601-L-CE, 1:250), CD163 (10D6) (Leica

Biosystems CD163-L-CE, 1:200), Anti-AREG (Sigma Aldrich HPA008720, 1:25). An Opal tyramide signal amplification system was used for fluorescent detection (Akoya Biosystems), and a DAPI nuclear counterstain was applied. The resulting assay was applied to 4 µm thick TMA sections. Whole slide images were captured at 10x magnification using the Vectra Polaris multispectral slide scanner (Akoya Biosystems), and individual core multispectral images were captured at 20x magnification. A spectral library was created using single stained slides of lung adenocarcinoma with each marker-fluorophore pair, and an autofluorescence control was created using an unstained section of lung adenocarcinoma. Images were spectrally unmixed using Inform software (Akoya Biosciences version 2.6.0). The generated component images were analysed using Visiopharm image analysis software (v2022.12.0.12865). A deep learning classifier was used to segment core images into background, tumour, stroma, and necrosis regions. An additional deep learning classifier used DAPI nuclear counterstain to detect cells within regions of interest. Mean intensity measures were generated for each marker at the cell and core level.

**SUPPLEMENTARY FIGURE LEGENDS**

**Figure S1: Characterisation of APOBEC3B expressing tumours**

**A)** Histological confirmation of liver metastasis stained with H&E. Scale bars = 100µm. T = tumour; L = normal liver. Quantification of visible metastasis (identified by eye or using PEARL imager) in KM (N = 5) and KMA (N = 6) mice induced with AS-CRE at $1x10^7$ pfu. **B)** Representative images of *in situ* hybridisation of human A3B in KM and KMA mice at 8-weeks post allele induction and at end stage. Scale bars = 100µm. **C)** Immunoblots of $KRAS^{G12D}$, MYC, Vinculin (top 3 rows all from the same gel), pan-RAS & Vinculin (from the same lysates run on a second gel) expression in lysates from primary end-stage lung tumours of KM and KMA mice (N = 4 biological replicates). Dashed line indicates removal of an empty lane from images.

### Figure S1

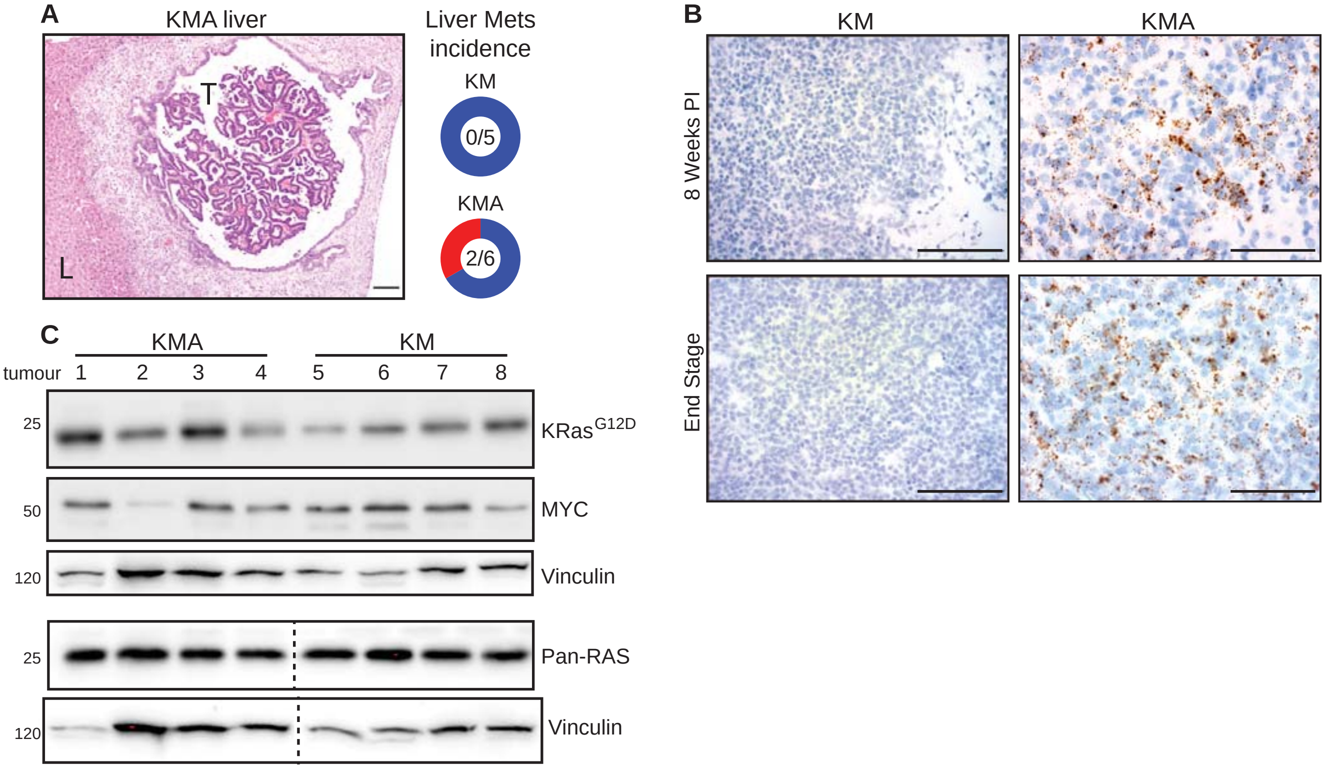
A
KMA liver
T
L
Liver Mets incidence
KM
0/5
KMA
2/6
B
KM
KMA
8 Weeks PI
End Stage
C
KMA
KM
tumour 1 2 3 4 5 6 7 8
25
KRas G12D
50
MYC
120
Vinculin
25
Pan-RAS
120
Vinculin

**Figure S2: APOBEC3B mutagenesis & Cytidine deaminase activity**

**A)** Absolute quantification of mutations in lung tumour samples, compared with matched normal liver, from KM (N = 6) and KMA (N = 15) mice harvested at clinical end point. COSMIC single base substitution signatures indicated by colour. **B)** Relative mutation burden per megabase in tumour samples from KM and KMA mice, as per (A). Ns = not significant (unpaired T test). **C)** Immunoblots of lysates generated from KP and KPA cell lines derived from primary tumours of the indicated genotype, using antibodies indicated. **D)** KPA cell lines demonstrate increased deaminase activity compared to KP cell lines. Deamination assay was quantified by densitometry of cleaved substrate using Image J (N = 3). **E)** Apurinic (AP) sites on DNA extracted from KP and KPA cell lines indicate an increase in the number of AP sites in KPA DNA compared with KP DNA (N = 3 repeat cultures of each cell line). **** denotes $p<0.0001$; ** denotes $p<0.01$, ns = not significant (ANOVA & post-hoc Tukey test)

### Figure S2

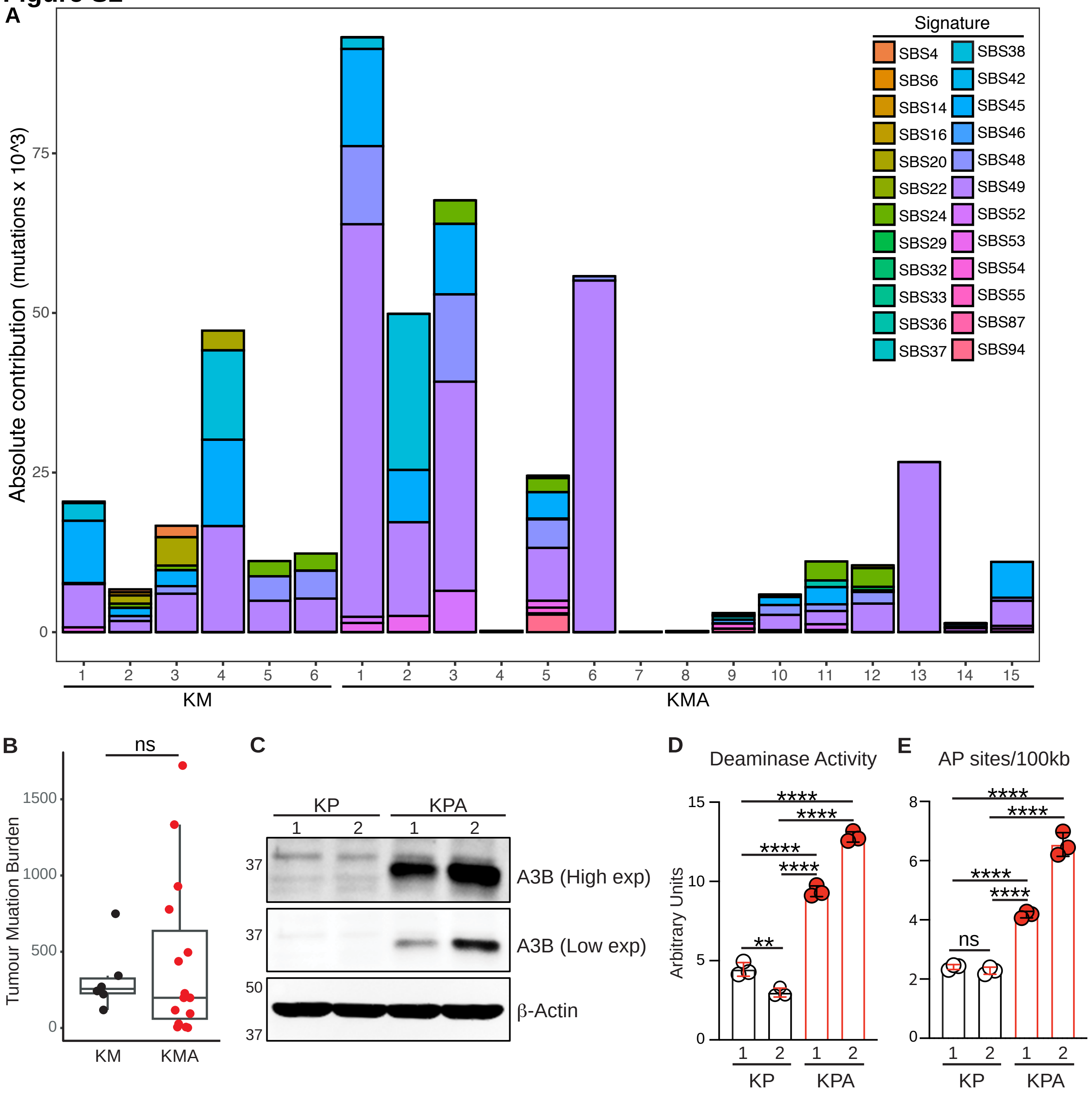

A
Signature
SBS4
SBS6
SBS14
SBS16
SBS20
SBS22
SBS24
SBS29
SBS32
SBS33
SBS36
SBS37
SBS38
SBS42
SBS45
SBS46
SBS48
SBS49
SBS52
SBS53
SBS54
SBS55
SBS87
SBS94
Absolute contribution (mutations x 10^3)
KM
KMA
B
ns
Tumour Muation Burden
KM
KMA
C
KP
KPA
A3B (High exp)
A3B (Low exp)
β-Actin
D
Deaminase Activity
Arbitrary Units
E
AP sites/100kb

**Figure S3: Immune related gene expression in KMA tumours**

All panels show normalised RNA-Seq reads from tumour bearing lungs of KM and KMA mice harvested at 8-weeks post allele induction (N = 4 biological replicates) of **A)** genes associated with IFN signalling; **B)** of chemokines and receptors associated with anti-tumour immunity; **C)** of IghM expression; and **D)** of genes associated with T cell function. *** denotes adjusted p<0.001; ** denotes adjusted p<0.01; * adjusted p<0.05 (unpaired T-test).

### Figure S3

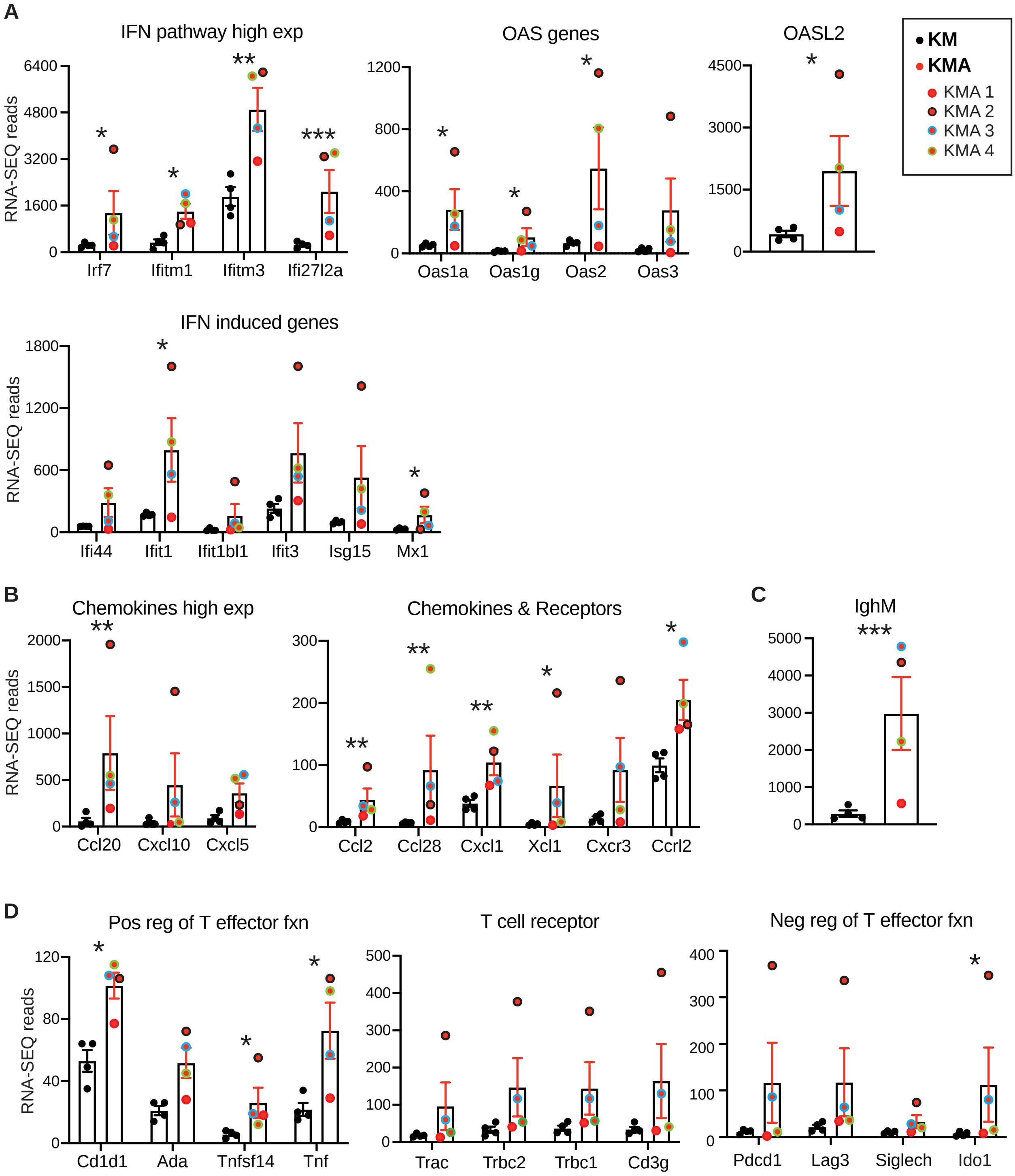

A
IFN pathway high exp
RNA-SEQ reads
6400
4800
3200
1600
0
Irf7
Ifitm1
Ifitm3
Ifi27l2a
OAS genes
1200
800
400
0
Oas1a
Oas1g
Oas2
Oas3
OASL2
4500
3000
1500
0
KM
KMA
KMA 1
KMA 2
KMA 3
KMA 4
IFN induced genes
1800
1200
600
0
Ifi44
Ifit1
Ifit1bl1
Ifit3
Isg15
Mx1
B
Chemokines high exp
2000
1500
1000
500
0
Ccl20
Cxcl10
Cxcl5
Chemokines & Receptors
300
200
100
0
Ccl2
Ccl28
Cxcl1
Xcl1
Cxcr3
Ccrl2
C
IghM
5000
4000
3000
2000
1000
0
D
Pos reg of T effector fxn
120
80
40
0
Cd1d1
Ada
Tnfsf14
Tnf
T cell receptor
500
400
300
200
100
0
Trac
Trbc2
Trbc1
Cd3g
Neg reg of T effector fxn
400
300
200
100
0
Pdcd1
Lag3
Siglech
Ido1

**Figure S4: Immune landscape of KM and KMA tumours**

**A)** Representative images of KM and KMA lungs at 8-weeks post allele induction stained for NKp46, CD45R, F4/80, and Ly6G. Scale bars = 100µm. **B)** HALO quantification of tumour infiltrating NK cells, B cells, macrophages, and neutrophils. Each symbol denotes mean value across 1 lung section from each mouse. Error bars represent Mean ± SD. * denotes $p<0.05$ (unpaired T-test). **C)** Representative images of CD45R staining of KM and KMA lungs at 8-weeks post allele induction at low magnification showing tertiary lymphoid structures forming around tumours and blood vessels in both KM and KMA tumour bearing lungs. Scale bars = 100µm. **D)** Correlation between tumour infiltrating CD8$\alpha^+$ T cells and the number of tertiary lymphoid structures in adjacent sections of lung tissue. N=7 KM and 8 KMA mice. The coefficient of determination ($R^2$) indicates a positive correlation between the variables in both genotypes (calculated in GraphPad). **E)** Representative images of end-stage lungs from KM and KMA mice stained for NKp46, CD45R, F4/80, and Ly6G. Scale bars = 100µm.

#### Figure S4

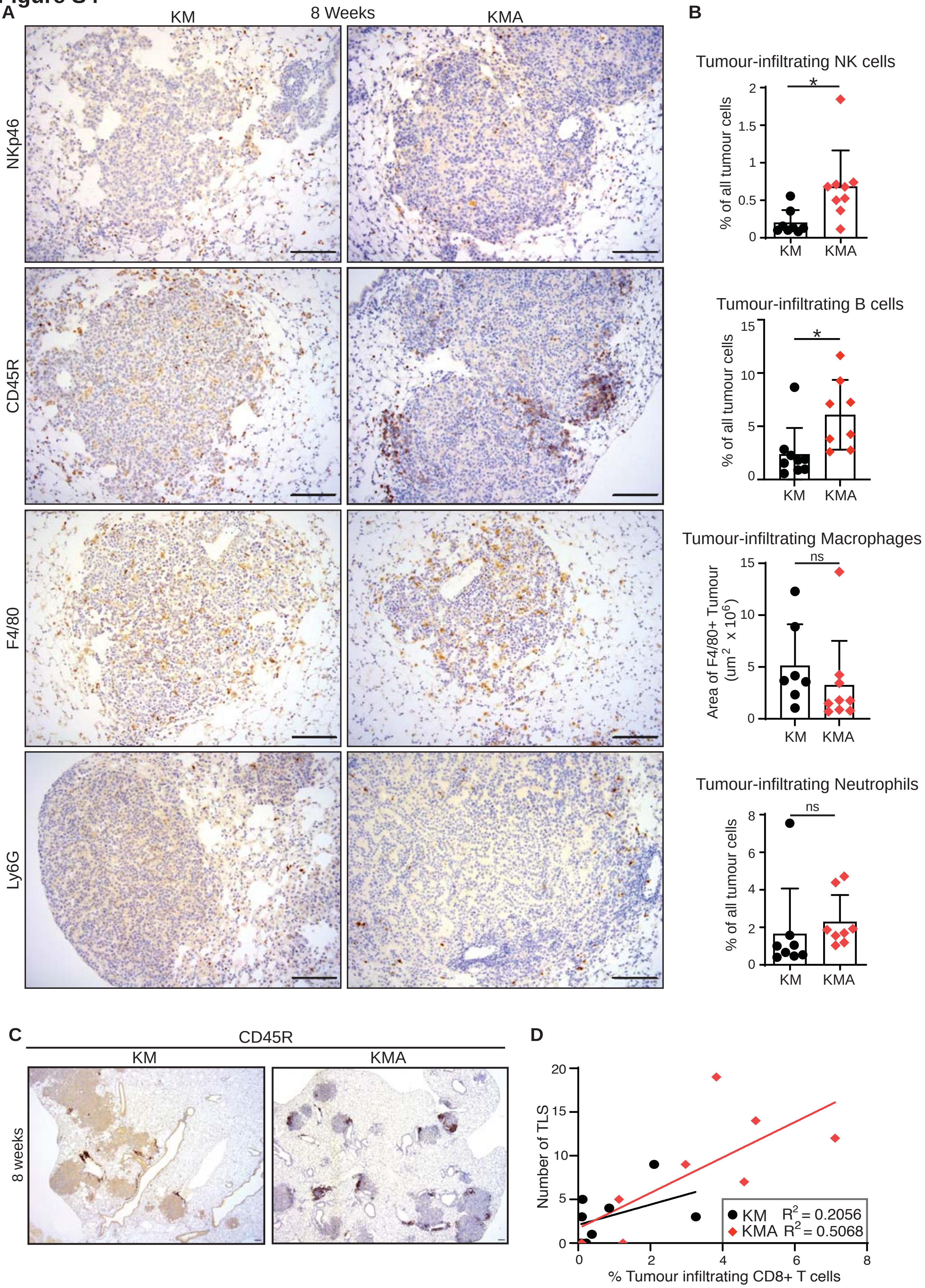

A
8 Weeks
KM
KMA
NKp46
CD45R
F4/80
Ly6G
B
Tumour-infiltrating NK cells
% of all tumour cells
*
KM
KMA
Tumour-infiltrating B cells
% of all tumour cells
*
KM
KMA
Tumour-infiltrating Macrophages
Area of F4/80+ Tumour (um2 x 10^6)
ns
KM
KMA
Tumour-infiltrating Neutrophils
% of all tumour cells
ns
KM
KMA
C
CD45R
KM
KMA
8 weeks
D
Number of TLS
% Tumour infiltrating CD8+ T cells
KM R2 = 0.2056
KMA R2 = 0.5068

**Figure S4 (continued)**

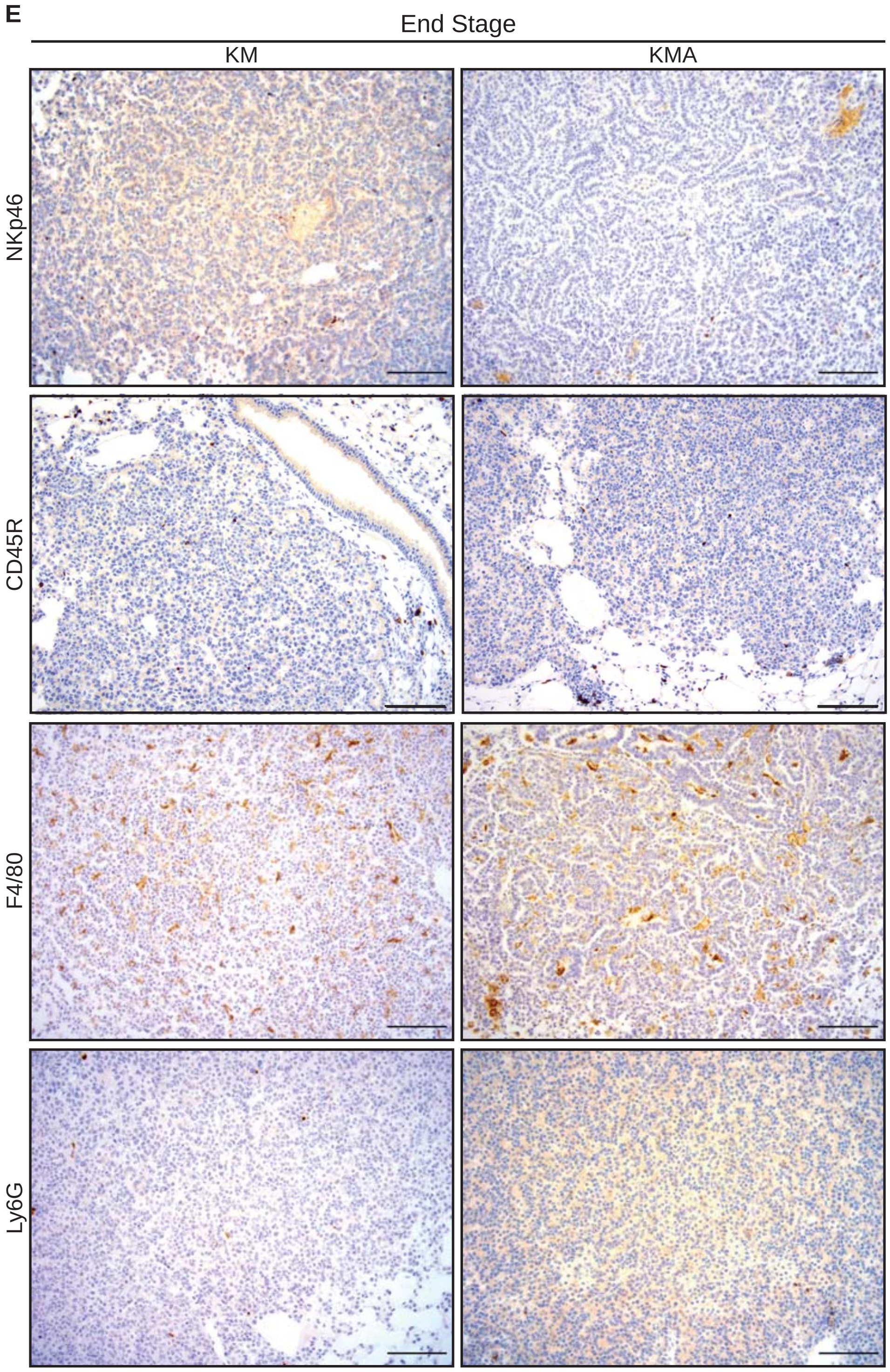

**Figure S5: Depletion of CD8 T cells negates KMA survival advantage**

**A)** Representative contour plots of $CD8^+$ T cells detected in blood samples from KM and KMA mice, 50-days post allele induction on day 0 prior to first treatment with CD8-depleting antibody, and at day 14 (x4 doses), measured by FACS. **B)** Quantification of circulating CD8+ T cells in KM and KMA mice on day 0 and day 14 of anti-CD8+ depleting antibody treatment. Error bars represent Mean ± SD (paired T-test). **C)** Overall survival of KM (N = 7) and KMA (N = 9) mice treated with anti-CD8 depleting antibody. Graph directly compares CD8-depelted KM (from main Figure 3C) and KMA (from main Figure 3B) mice. Mantel-Cox log rank test. For all panels, ns denotes not significant; **** denotes p <0.0001.

#### Figure S5

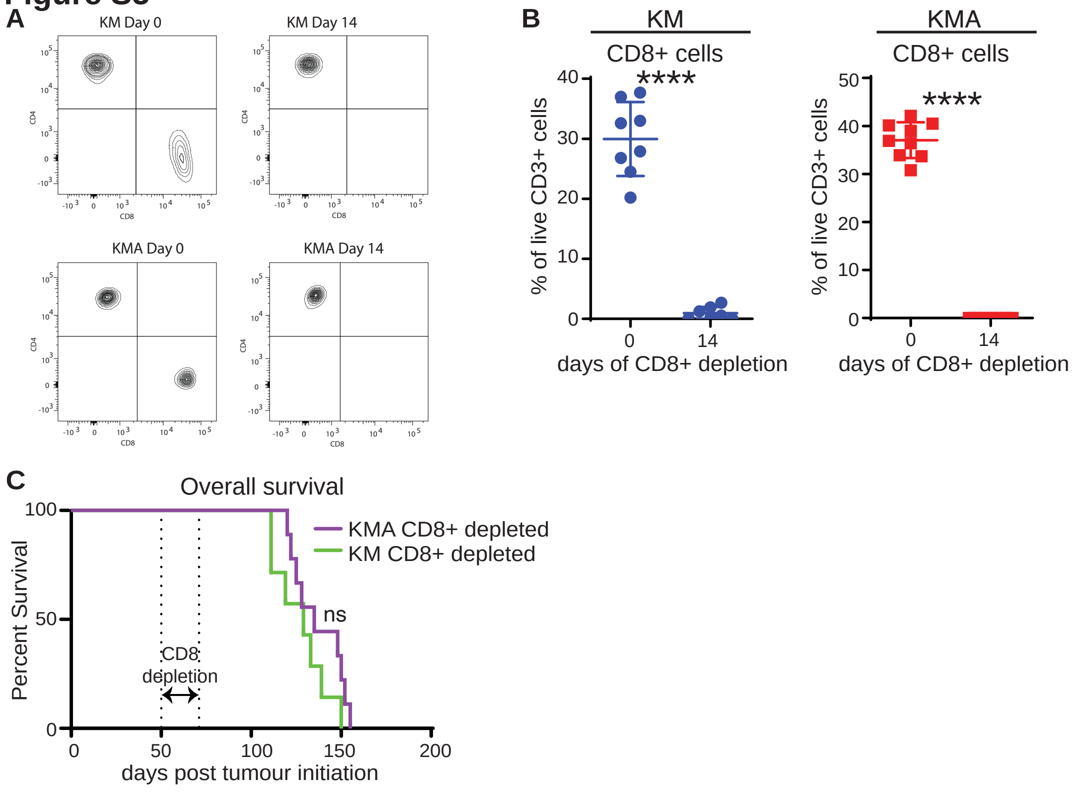

A
KM Day 0
KM Day 14
KMA Day 0
KMA Day 14
CD4
CD8
B
KM
CD8+ cells
KMA
CD8+ cells
% of live CD3+ cells
****
days of CD8+ depletion
C
Overall survival
Percent Survival
KMA CD8+ depleted
KM CD8+ depleted
ns
CD8 depletion
days post tumour initiation

**Figure S6: Gene expression changes induced by inhibition of the MYC repressor complex**

**A)** Representative CD4 IHC of a super responder KMA tumour following treatment with 1mg/kg Trametinib daily for 5 days immediately prior to harvest at 12-weeks post allele induction (day 78 to day 83). Scale bars = 100μM. **B)** Representative CD4 staining of super responder KMA tumour following treatment with 40mg/kg HKMTI-1-005, as per (A). Scale bars = 100μM. **C)** Schematic of pathway enrichment from tumour bearing lungs of KMA mice treated with 40mg/kg HKMTI-1-005 or vehicle control twice daily for 5 days immediately prior to harvest at 12-weeks post allele induction and analysed by RNA-Seq. The top 12 significantly up-regulated and down-regulated pathways were identified using Metacore GeneGO analysis. **D)** Normalised RNA-Seq reads of indicated Interferon-responsive genes in tumour-bearing lungs from KMA mice treated with HKMTI-1-005 (N = 4) or vehicle (N = 4) as per (C). Mean and SEM shown. **E)** Normalised RNA-Seq reads of indicated B cell maturation and class identifier genes in KMA mice treated with HKMTI-1-005 or vehicle as per (C). **F)** Representative IHC of IgG expression in tumour-bearing lungs of KMA mice treated with 40mg/kg HKMTI-1-005 (N = 3) or vehicle control (N = 3) twice daily for 5 days immediately prior to harvest at 12-weeks post allele induction. For all panels *** denotes adjusted $p<0.001$; ** $p<0.01$; * $p<0.05$ (unpaired T-test).

### Figure S6

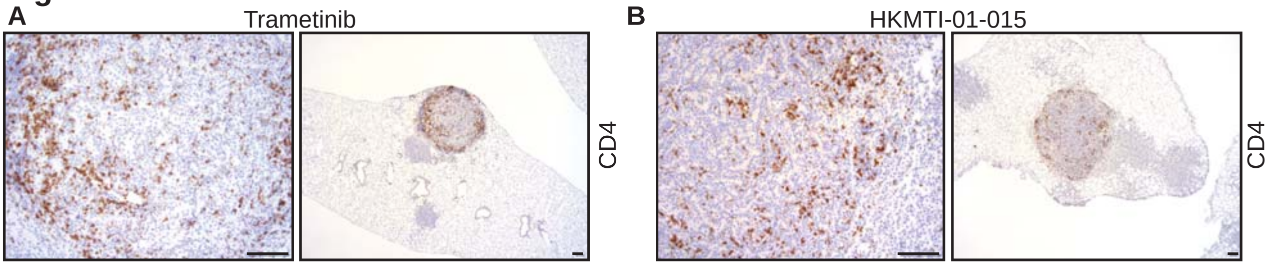


C

| Up-regulated pathways | FDR |
|---|---|
| 1 Immune response: IFNα/β signalling via JAK/STAT | 1.12e-7 |
| 2 Immune response: Alternative Complement pathway | 1.12e-7 |
| 3 Alternative complement cascade in age-realted macular degeneration | 4.99e-7 |
| 4 Immune response: Antigen presentation by MHC class I: cross-presentation | 2.19e-5 |
| 5 Cell adhesion: Integren inside-out signalling in neutrophils | 5.08e-5 |
| 6 Immune response: Antiviral actions of interferons | 5.74e-5 |
| 7 Immune response: IFNa/b signalling via MAPKs | 1.65e-4 |
| 8 Chemotaxis SDF-1/CXCR4-induced chemotaxis of immune cells | 2.91e-4 |
| 9 Impaired macrophage phagocytic function in asthma | 2.91e-4 |
| 10 COVID-19 associated coagulopathy | 2.91e-4 |
| 11 Complement pathway disrupton in thrombotic microangiopathy | 2.93e-4 |
| 12 SLE genetic marker-specific pathways in antigen-presenting cells (APC) | 3.99e-4 |

| Down-regulated pathways | FDR |
|---|---|
| 1 Cell cycle: The metaphase checkpoint | 4.32e-30 |
| 2 Cell cycle: Chromosome condensation in prometaphase | 1.16e-22 |
| 3 Cell cycle: Role of APC in cell cycle regulation | 1.94e-18 |
| 4 DNA damage: ATM/ATR regulation of G2/M checkpoint (nuclear signalling) | 1.29e-15 |
| 5 DNA damage: ATM/ATR regulation of G2/M checkpoint (cytoplasmic signalling) | 7.63e-15 |
| 6 DNA damage: Intra S-phase checkpoint | 9.18e-12 |
| 7 Cell cycle: Inititation of mitosis | 1.16e-11 |
| 8 Cell cycle: Spindle assmebly and chromosome separation | 2.29e-10 |
| 9 DNA damage: Double-strand break repair via homologous recombination | 4.74e-10 |
| 10 Cell cycle: Start of DNA replication in early S-phase | 3.86e-9 |
| 11 Cell cycle: Role of Nek in cell cycle regulation | 9.43e-8 |
| 12 Cell cycle: Transition and termination of DNA replication | 3.23e-7 |

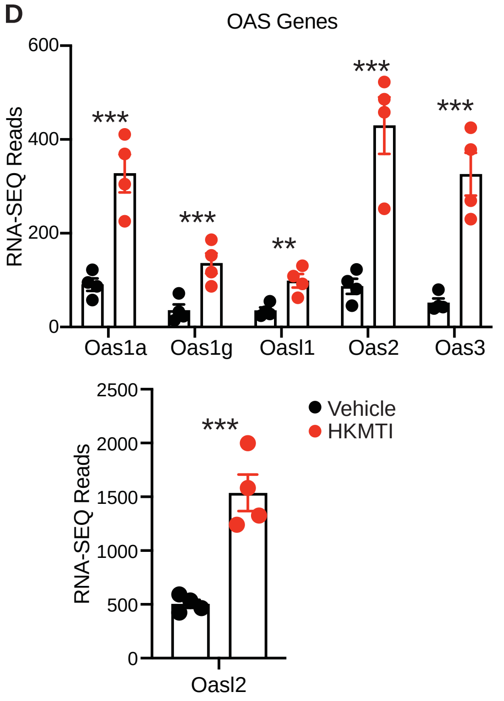


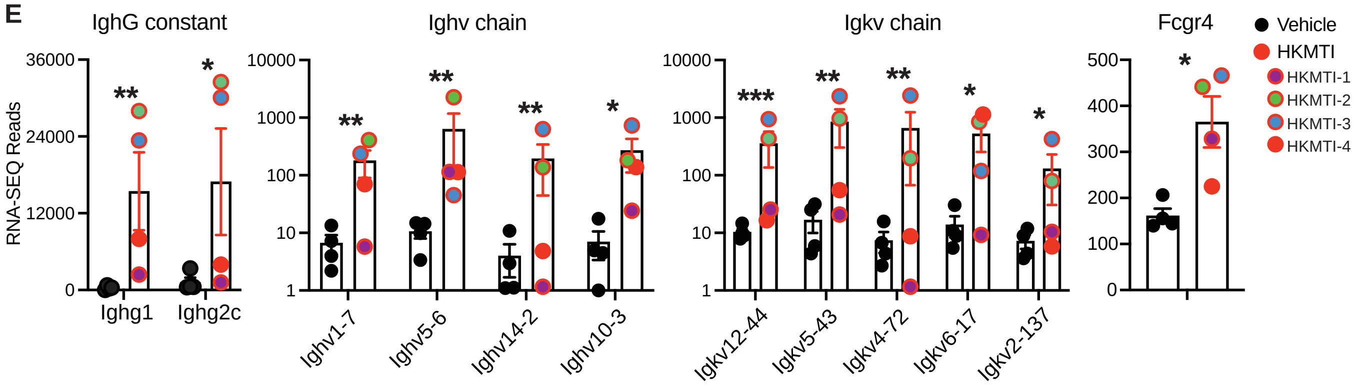


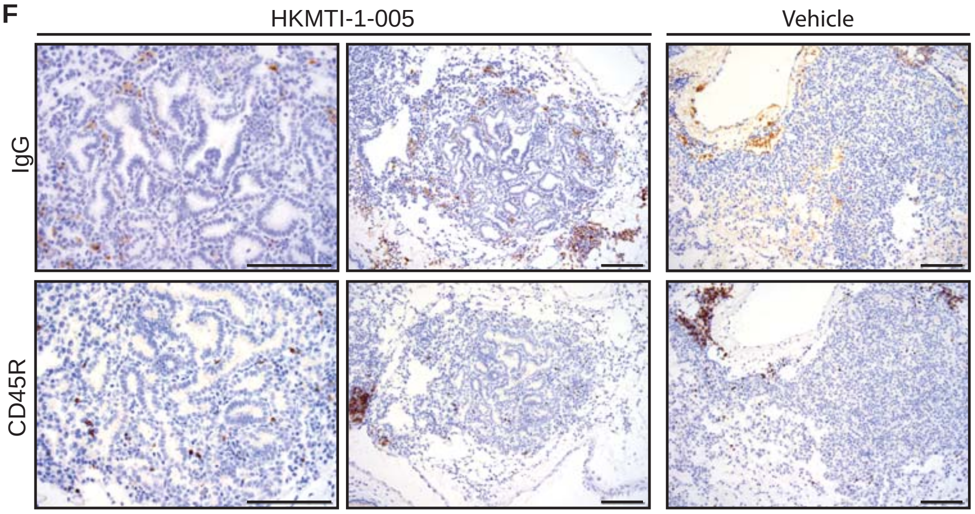

**SUPPLEMENTARY TABLE LEGENDS**

**Table S1: Cell Lineage Assignments**
Genes associated with each cluster and cell lineage identity, as determined using the Immgen online identifier tool (see supplementary methods below).

**Table S2: Pathway Enrichment**
Gene ontology (GO) biological process comparison of clusters 3 & 12, performed using PANTHER overrepresentation analysis
